# Egocentric and Allocentric Representations in Auditory Cortex

**DOI:** 10.1101/098525

**Authors:** Stephen M. Town, W. Owen Brimijoin, Jennifer K. Bizley

## Abstract

A key function of the brain is to provide a stable representation of an object’s location in the world. In hearing, sound azimuth and elevation are encoded by neurons throughout the auditory system and auditory cortex is necessary for sound localization. However the coordinate frame in which neurons represent sound space remains undefined: classical spatial receptive fields in head-fixed subjects can be explained either by sensitivity to sound source location relative to the head (egocentric) or relative to the world (allocentric encoding). This coordinate frame ambiguity can be resolved by studying freely moving subjects and here we recorded spatial receptive fields in auditory cortex freely moving ferrets. We found two distinct populations of neurons: While the majority (∼80%) of spatially tuned neurons represented sound source location relative to the head, we provide novel evidence for a group of neurons in which space was represented in an allocentric world-centered coordinate frame. We also use our ability to measure spatial tuning in moving subjects to explore the influence of sound source distance and speed of head movements on auditory cortical activity and spatial tuning. Modulation depth of spatial tuning increased with distance for egocentric but not allocentric units, whereas for both populations modulation was stronger at faster movement speeds. Our findings argue that auditory cortex is involved in the representation of both sound source location relative to ourselves and sound location in the world independent of our own position.

## Introduction

A key function of the brain is to provide a stable representation of an object’s location in the world. In hearing, this requires an observer maintains the identification of an auditory object as they move through an environment. Movement is a critical aspect of sensing [1] that contributes to sound localization and other auditory behaviors [2-7], however the neural basis underpinning active hearing and world centered auditory representations remains unknown.

For a moving observer, it is possible to represent sound location either relative to oneself (egocentric representation) or relative to the world through which one moves (allocentric representation). Allocentric representations provide a consistent report of object location across movement of an observer [8], as well as a common reference frame for mapping information across several observers or multiple sensory systems [9, 10]. Despite the computational value and perceptual relevance of allocentric representations to hearing, studies of auditory processing have only recently considered the coordinate frames in which sound location is represented [11-13]. Both electroencephalography (EEG) and modelling studies hint that sound location might be represented in cortex in both head-centered and head-independent spaces. However EEG has not yet revealed the precise location of these representations and cannot determine how individual neurons in tonotopic auditory cortex define space.

In static subjects, auditory cortical neurons encode sound azimuth and elevation [14-18] and localization of sound sources requires an intact auditory cortex [19-21]. However, in static subjects with a fixed head position, neural tuning to sound location is ambiguous, as the head and world coordinate frames are fixed in alignment and so allocentric and egocentric sound location are always equal. While it has been largely assumed cortical neurons represent sound location relative to the head, the spatial coordinate frame in which location is encoded remains to be demonstrated. Furthermore, though the acoustic cues to sound localisation are explicitly head-centered, information about head direction necessary to form a world-centered representation is present at early levels of the ascending auditory system [22]. Thus it may be possible for neurons in the auditory system to represent space in an allocentric, world-centered coordinate frame that would preserve sound location across self-generated movement.

Here we resolve the coordinate frame ambiguity of spatial tuning in auditory cortex by recording from neurons in freely moving subjects. In moving conditions, the head and world coordinate frames are no longer fixed and so we can determine in which coordinate frame a given cell is most sensitive. Our approach reveals distinct populations of head-centered and world-centered cells that suggest the coexistence of egocentric and allocentric representations in auditory cortex. We also explore the impact that distance from a sound source and subject’s self-generated movement have on these distinct neuronal sub-populations.

## Results

We hypothesised that measuring spatial tuning in moving subjects would allow us to distinguish between egocentric (head-centered) and allocentric (world-centered) representations of sound location (Fig 1). To formalize this theory and develop quantitative predictions about the effects of observer movement on spatial tuning, we first simulated egocentric and allocentric neurons that were tuned to sound locations defined relative to the head (Fig 1a) and world (independent of the subject) respectively (Fig 1b). For both these units, we identified parameters that produced the same spatial tuning in response to sounds presented from a classical speaker ring with the head at the center (Fig 1c-d). The overlap in tuning obtained from egocentric and allocentric representations numerically illustrates coordinate frame ambiguity. However our simulation confirmed that when the observer moved freely with a uniform distribution of head directions, spatial tuning would only be apparent in the coordinate frame relevant for neural output (Fig 1e). Additionally, changes in head direction would produce systematic shifts in tuning curves in the coordinate frame that was irrelevant for neural output while tuning in the relevant coordinate frame would be invariant across head direction (Fig 1f). We subsequently demonstrated that tuning curves of many shapes and preferred locations can theoretically be explained by spatial receptive fields based within an allocentric coordinate frame (Supplementary Fig. 1). With simulations providing a foundation, we then made recordings in freely moving animals to determine whether the spatial tuning of auditory cortical neurons followed egocentric or allocentric predictions.

**Fig 1.**
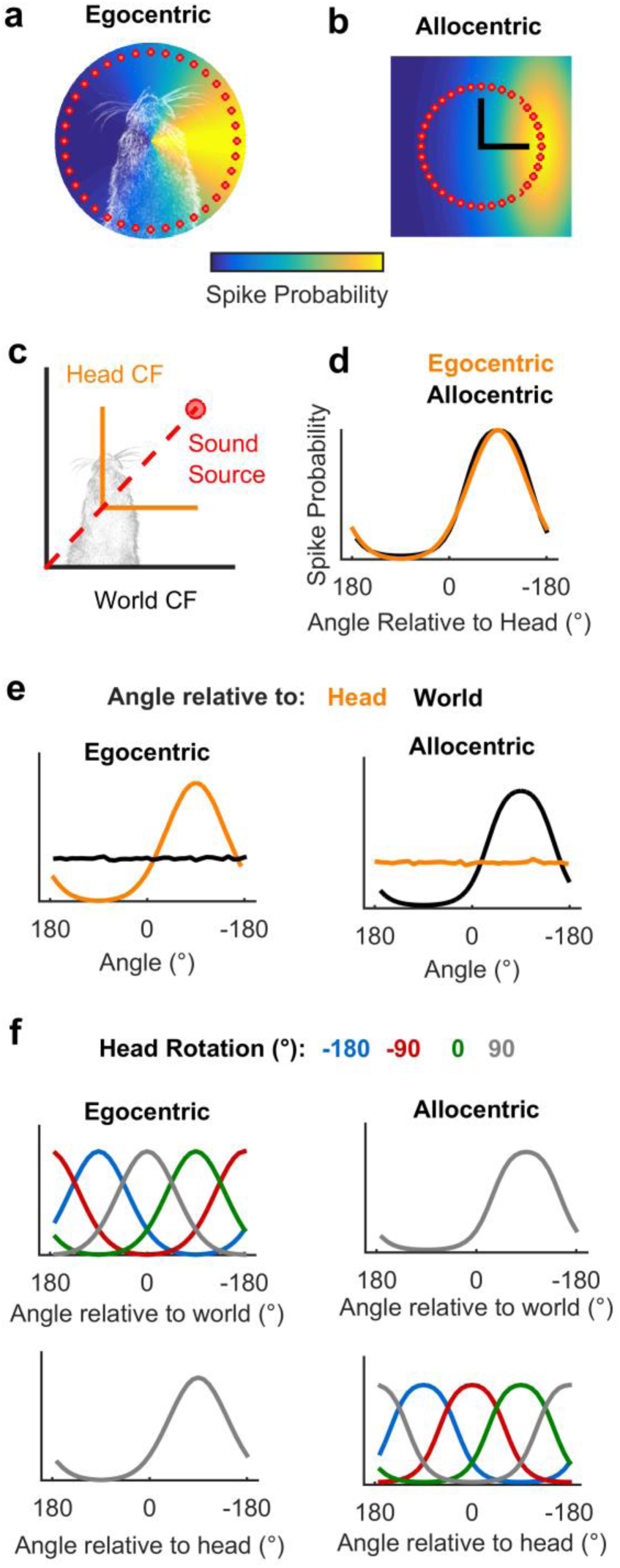
Simulated receptive fields show that observer movement resolves coordinate frame ambiguity. **a-b**, Simulated neurons with receptive fields tuned to sound location relative to the head (a, Egocentric) or in the world (b, Allocentric). Circles show hypothetical sound sources in a classical speaker ring; black lines indicate axes and origin of the simulated world. **c**, Schematic of world and head coordinate frames (CF). **d**, Sound-evoked tuning curves according to allocentric and egocentric hypotheses when head and world coordinate frames aligned. **e**-**f**, predictions of allocentric and egocentric hypotheses across head rotation and movement (e) and at specific head directions (f).

To measure spatial tuning in moving subjects, we implanted ferrets (n = 5) with multi-channel tungsten electrode arrays allowing the recording of single and multi-unit activity during behavior. During neural recording, ferrets were placed in an arena which the animals explored for water rewards while the surrounding speakers played click sounds (Fig 2a). To measure the animal’s head position, direction and speed in the world during exploration (Fig. 2b-f) we tracked LEDs placed on the midline of the head (Supplementary Video 1). During exploration, click sounds were presented from speakers arranged at 30° intervals between ± 90° relative to the arena center with speaker order and inter-click interval varied pseudo randomly. We also roved the level of clicks between 54 and 60 dB SPL such that absolute sound level varied both as a function of sound source level and distance between head and speaker, and absolute sound level did not provide a cue to sound location (Fig 2g-h). Clicks were used as they provided instantaneous energy and thus ensured minimal movement of the animal during stimulus presentation (Supplementary Fig. 3c-d). The locations (speaker angle) from which the clicks originated were used alone to estimate *allocentric* receptive fields and were used in conjunction with the animal’s head direction and position to deconvolve *egocentric* spatial receptive fields.

**Fig 2.**
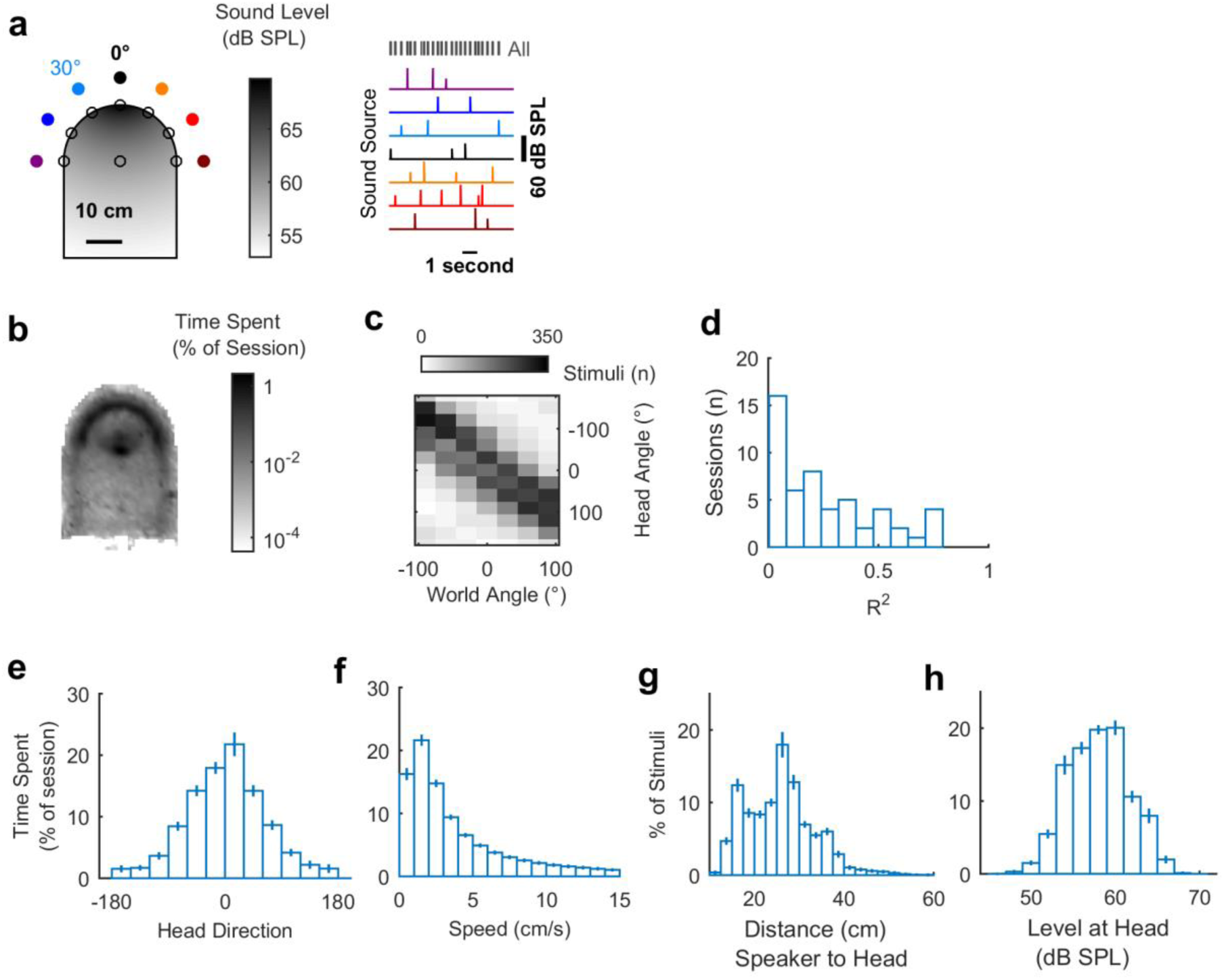
Experimental design and exploratory behavior in a sound field. **a**, Arena with speakers (filled circles) and water ports (unfilled circles). Shading indicates the sound field generated by a click from the speaker at 0° calibrated to be 60 dB SPL at the center of the chamber. Stimuli were presented with a pseudorandom interval and order across speakers. **b**, Mean proportion of time in each recording session spent within the arena. **c**, stimulus angles relative to the head and world for one session that was representative of behavior in all sessions (n = 57). **d**, Correlation coefficients (R^2^) between sound angles in head and world coordinate frames across all behavioral sessions. **e-h**, Distributions of head direction, head speed, distance between head and sound source and the sound level at the animal’s head during behavior. Bars indicate mean ± s.e.m. across sessions.

We observed that animals moved throughout the arena to collect water (Fig 2b) and used a range of head directions during exploration (Fig 2e). In contrast to our initial simulations, the distribution of the animal’s head directions was notably non-uniform, leading to correlations between sound source angle relative to the head and the world (e.g. Fig 2c; mean ± s.e.m. R^2^ = 0.247 ± 0.0311). This correlation between sound source angles resulted because the animal preferred to orient towards the midline of the arena and thus sounds that were to the right of the animal were more often on the right of the arena than would result from random behavior. The preference of the animal was likely a consequence of the shape of the arena, and although relatively small, we sought to determine how the animal’s head direction preference affected our experimental predictions.

To assess the influence of real animal behavior on our ability to distinguish coordinate frames, we combined the head directions observed during behavior with simulated neural receptive fields. We combined our simulated egocentric and allocentric receptive fields (Fig 1) with the animal’s head position and direction across each single behavioural testing session (Fig 3a) to calculate the spatial tuning for allocentric and egocentric units in both head and world coordinate frames. Our model predictions (Fig.1) demonstrated that for a uniform distribution of head angles, the tuning function of allocentric or egocentric unit should be flat when considered in the irrelevant coordinate frame. A bias in head location over time however, would produce modulation in firing rate with location in the irrelevant coordinate frame, appearing as tuning in that coordinate frame (Fig 3a). In order to account for this we therefore measured the modulation depth of each tuning curve and calculated the strength of the *residual modulation:* the modulation depth in the simulation-irrelevant coordinate frame (e.g. in the world coordinate frame for an egocentric unit) expressed as a percentage of modulation depth in the simulation-tuned coordinate frame (e.g. the head coordinate frame for an egocentric unit)( Fig. 3, see also Methods, Equation 11). Residual modulation thus represents the degree of indirect spatial tuning in one coordinate frame observed as a by-product of spatial tuning in another coordinate frame combined with the animal’s behavior.

**Fig 3.**
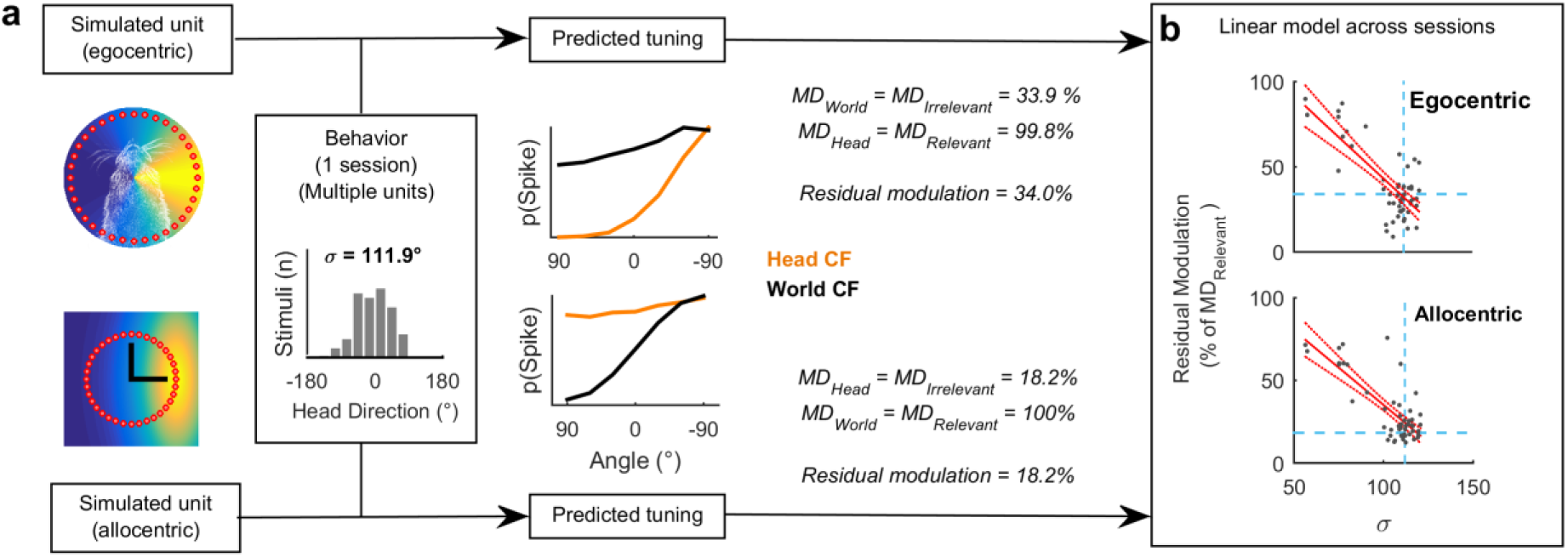
Estimating residual modulation. **a**, Example workflow for estimating residual modulation in coordinate frames irrelevant for neural output that result from biases in head direction. Estimations performed separately for each behavioral session. **b** Inverse correlation between residual modulation standard deviation of head directions (σ). Red filled lines indicate regression fit and confidence intervals. Dashed lines indicate data point for the single session in (a).

Across all behavioral sessions, residual modulation was inversely correlated with variation in the animal’s head direction (expressed as standard deviation) for both egocentric (R^2^ = 0.562, *p* = 1.07 x 10^-10^) and allocentric simulated units (R^2^ = 0.615, *p* = 3.73 x 10^-12^) (Fig 3b). This indicated that for real animal behavior, we would not expect to see the complete abolition of tuning but rather spatial tuning in both coordinate frames, with the weaker spatial tuning potentially attributable to the animal’s bias in head direction. In our neural analysis, we thus used the relationship between behavior and residual modulation to provide a statistical framework in which to assess the significance of spatial modulation of real neurons.

### Egocentric and allocentric tuning in auditory cortex

During exploration we recorded responses of 186 sound-responsive units (50 single units, 136 multi-units) in auditory cortex (Supplementary Fig. 2). Electrode arrays were targeted to span the low frequency areas where the middle and posterior ectosylvian gyral regions meet and thus units were sampled from primary auditory cortex and two tonotopically organised secondary fields: the posterior pseudosylvian and posterior suprasylvian fields. We analysed the firing rates of units in the 50 milliseconds after the onset of each click; this window was wide enough to capture the neural response while being sufficiently short that the animals head moved less than 1 cm (median 4 mm, Supplementary Fig. 3) and less than 30° (median 12.6°) – the interval between speakers. The interval between stimuli always exceeded 250 ms.

We identified periods of time when the animal was facing forwards at the center of the speaker ring: in this situation we mimic classic neurophysiological investigations of spatial tuning in which head and world coordinate frames are aligned. In the aligned case, 92 units that were significantly modulated by sound source location (Fig 4a-b and Supplementary Fig. 4) and spatial tuning curves computed in head and world coordinate frames were highly correlated (mean ± s.e.m. R^2^ = 0.889 ± 0.0131). We then compared the aligned control condition with all data when the head and world coordinate frames were free to vary. Compared to the aligned condition, correlation between egocentric and allocentric tuning curves was significantly reduced (Fig 4c, R^2^ = 0.522 ± 0.0294) (paired t-test: free vs. aligned *t*_91_ = 8.76, *p* < 0.001) and differences in spatial tuning in head and world coordinate frames became visible (Fig 4d and 4f).

**Fig 4.**
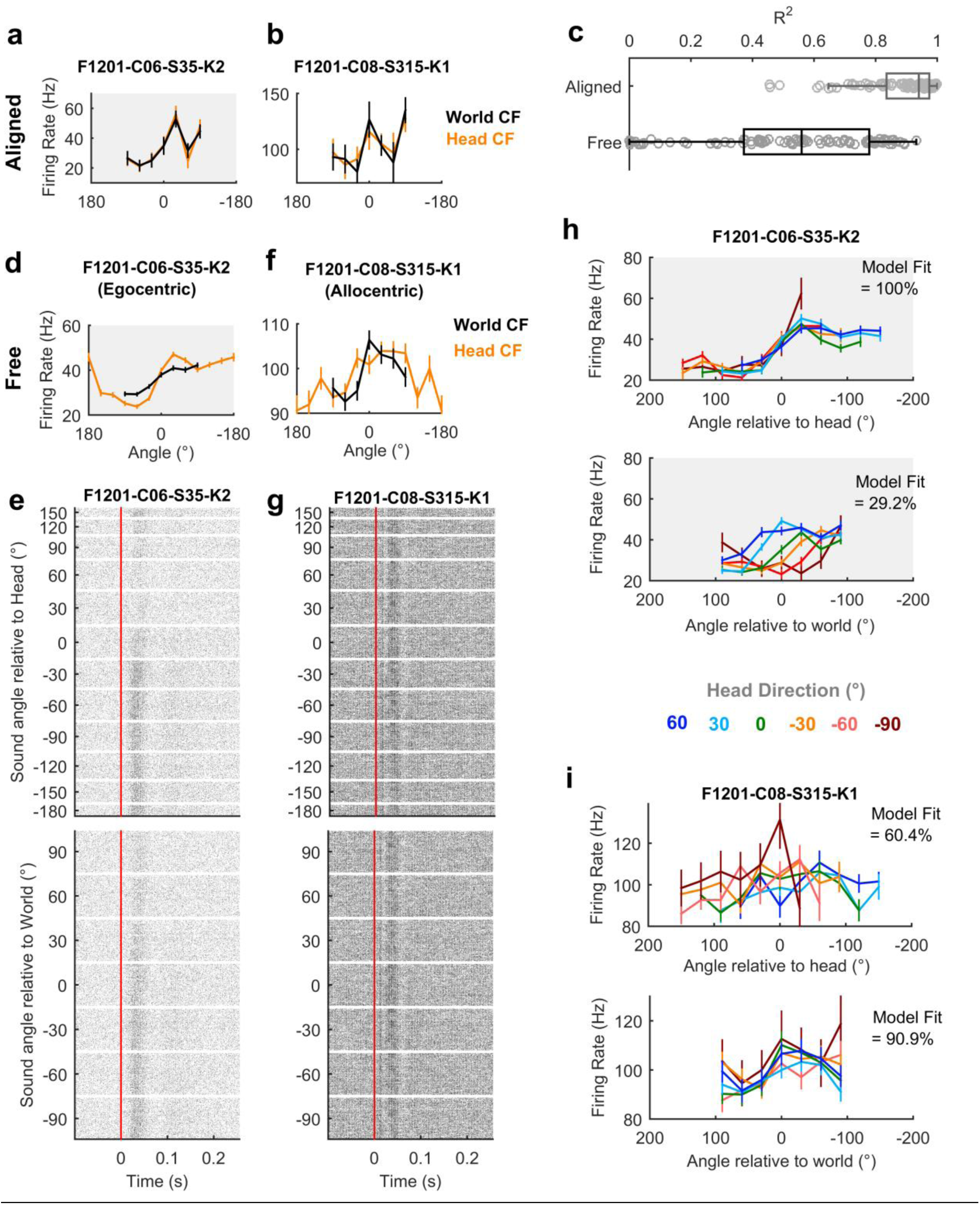
Spatial tuning of egocentric and allocentric units. **a-b**, Two simultaneously recorded units with spatial tuning curves matched when head and world coordinate frames (CF) are aligned. **c,** Correlation coefficient between tuning curves in head and world CFs when head and world CFs are aligned or free to vary as the animal foraged around the arena. **d**, Tuning curve of unit (same unit as in 4a) consistent with the egocentric hypothesis when considering all data in which the head and world coordinate frames were free to vary. **e**, Raster plots (n = 9963 stimulus presentations) of spike times for all responses plotted according to sound angle in head and world coordinate frames. **f-g**, Tuning curve (f) and raster plots (g) for a unit consistent with the allocentric hypothesis when considering all data when coordinate frames were free to vary. Example units were recorded during the same test sessions and thus trial structure of raster plots is preserved across units. **h**, Tuning curves for unit consistent with the egocentric hypothesis plotted across head rotations (60°, n = 1502 stimuli; 30°, n = 2339; 0°, n = 1658; −30°, n = 1750; −60°, n = 1096; −90°, n = 322;). Model fit refers to percentage of explainable deviance calculated according to Figure 6 across all data in which coordinate frames were free to vary. **i**, Tuning curves for unit consistent with allocentric hypothesis plotted across head directions (sample sizes same as for 4h). Data for tuning curves shown as mean ± s.e.m. firing rates.

When animals moved freely through the arena and head and world coordinate frames were thus dissociated, we observed units consistent with egocentric (Fig 4d-e and Supplementary Fig. 5) and allocentric hypotheses (Fig 4f-g and Supplementary Fig. 6). For units consistent with the egocentric hypothesis, spatial receptive fields were more strongly modulated by sound angle in the head than world coordinate frame (Fig 4d-e). Furthermore tuning curves for sounds plotted relative to the head remained consistent across head rotation but shifted systematically when plotted relative to the world (Fig 4h). Both outcomes are highly consistent with our simulation predictions (Fig 1).

In addition to identifying head-centered spatial tuning across movement, we also found units with spatial tuning that realized the predictions generated by the allocentric hypothesis. In contrast to the putative egocentric units, units consistent with the allocentric hypothesis showed greater modulation depth to sound angle in the world than head coordinate frame during free movement (Fig 4f-g and Supplementary Fig. 6). For allocentric units, spatial tuning in the world coordinate frame was robust to head rotation whereas tuning curves expressed relative to the head were systematically shifted when mapped according to head direction (Fig 4i and Supplementary Fig. 6).

### Modulation depth across coordinate frames

To quantify the observations we made above and systematically compare spatial tuning in world and head coordinate frames, we calculated modulation depth for both tuning curves for each unit (Fig. 5a). A key prediction from our simulations with both uniform head-directions (Fig.1) and actual head-directions (Fig 3a) was that modulation depth would be greater in the coordinate frame that was relevant for neural activity than the irrelevant coordinate frame (i.e. Head > World for egocentric; World > Head for allocentric). For 76 / 92 units (82.6%), we observed greater modulation depth in the head than world coordinate frame indicating a predominance of egocentric units.

**Fig 5.**
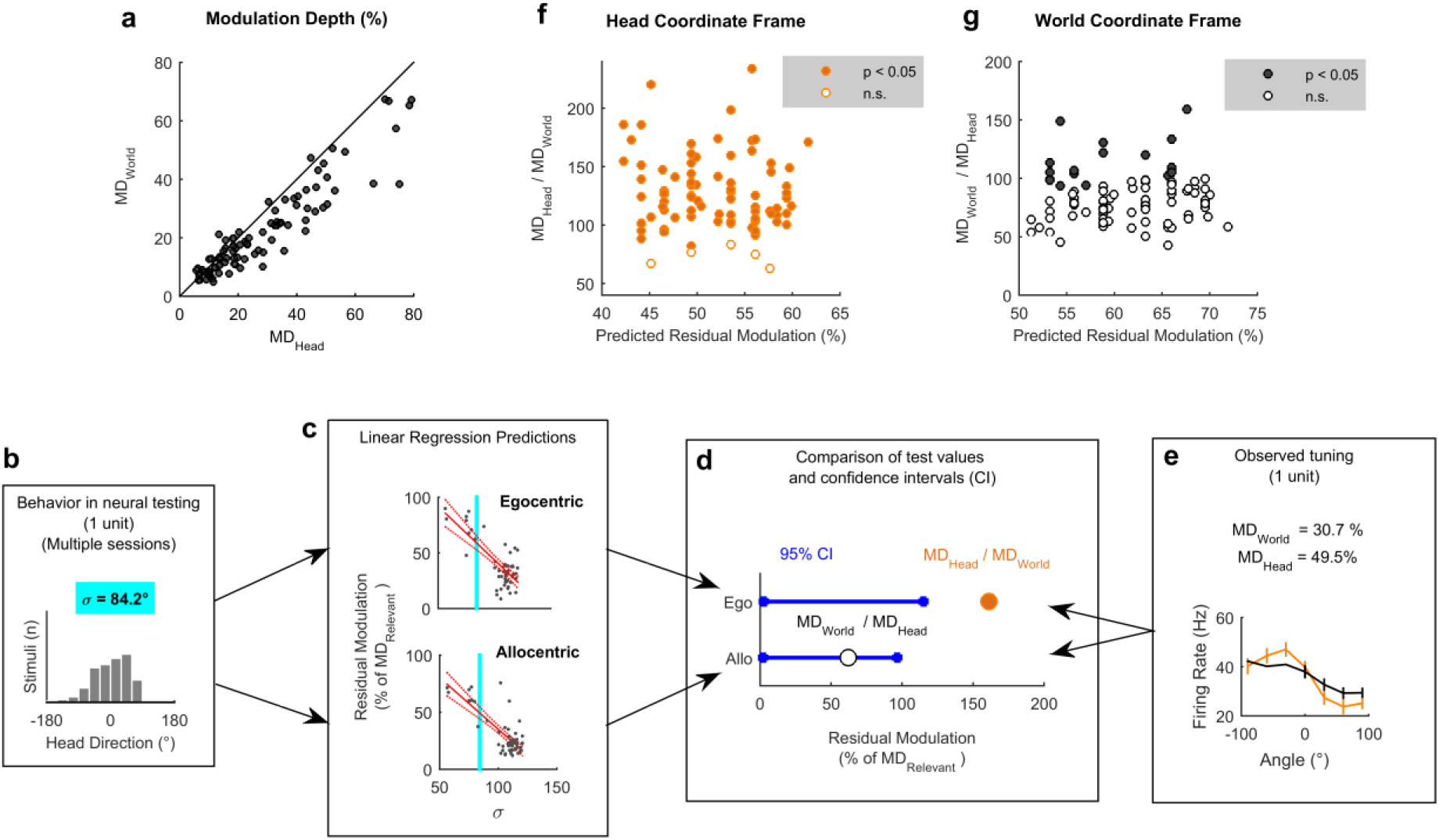
Modulation depth across coordinate frames. **a**, Modulation depth for all spatially modulated units (n = 92) compared in world (MD_World_) and head coordinate frames (MD_Head_) during exploration. **b-e** Workflow illustrating the use of animal behavior (b: summarized using the standard deviation of head directions during neural testing, σ) and linear regression models (c – see also Fig 3) to generate confidence intervals for residual modulation (d) that were compared to observed modulation depth values (e) normalized relative to the alternative coordinate frame. **f-g,** Normalized modulation depth observed for each spatially tuned unit compared against the mean residual modulation predicted from behavior in head (f) or world (g) coordinate frames.

We next asked if modulation depth observed in head and world coordinate frames was greater than the residual modulation predicted by our earlier simulations (Fig.3b). A linear regression model, developed using simulated receptive fields, was used to predict the magnitude of residual tuning for each coordinate frame from the animal’s behavior during recording of each unit (Fig 5b-e). To describe the animal’s behavior across the relevant behavioural testing sessions for each neural recording, we calculated the standard deviation of head directions (Fig. 5b). A smaller standard deviation indicates a less uniform range of head-directions and when combined with our regression model (Fig 5c) would predict higher residual modulation in both coordinate frames. Thus for a given standard deviation, we could use linear regression to obtain a predicted 95% confidence interval for the residual modulation in head and world coordinate frames arising from allocentric or egocentric tuning respectively (Fig. 5d). Test values of observed modulation were calculated by normalizing modulation depth in one coordinate frame by the other coordinate frame (Fig. 5e) and significance was attributed when test values exceeded the confidence interval (p < 0.05) of residual modulation.

Across all spatially tuned units, modulation in the head coordinate frame was significantly greater than the predicted residual modulation for 87 / 92 units (94.6%, Fig 5f); modulation in the world coordinate frame was significant for 19 / 92 units (20.7%, Fig 5g) which included all five of the units for which head-centered modulation was not significant. Thus our results suggest a population predominated by egocentric units but with a small population of allocentric units with distinct modulation characteristics and a further number of units (14 / 92) in which modulation depth was significantly greater than expected in both coordinate frames.

### General Linear Modelling to define egocentric and allocentric populations

Our analysis of modulation depth indicated the presence of both egocentric and allocentric representations in auditory cortex but also highlighted that the analysis of modulation depth alone was sometimes unable to resolve the coordinate frame in which units encoded sound location. However, to calculate modulation depth requires we discretize sound location into distinct angular bins and ignore single trial variation in firing rates by averaging neural responses across trials. General linear models (GLMs) potentially offer a more sensitive method as they permit the analysis of single trial data and allow us to treat sound angle as a continuous variable. We considered two models which either characterized neural activity as a function of sound source angle relative to the head (GLM_HEAD_) or in the world (GLM_world_). For all units for which at least one GLM provided a statistically significant fit (relative to a constant model, analysis of deviance; p < 0.05, 91 / 92 units), we compared model performance using the Akaike information criterion (AIC) for model selection. In accordance with the modulation depth analysis, the majority of units were better modelled by sound angle relative to the head than world (72/91 units; 79.1%; four animals) consistent with egocentric tuning. However, we also observed a smaller population of units (19/91 units; 20.9%; three animals) whose responses were better modelled by sound angle in the world and therefore represent space in allocentric coordinates.

To visualize GLM performance and explore egocentric and allocentric subpopulations further, we plotted a normalized metric of the deviance value usually used to assess model fit. Here we defined *model fit* as the proportion of explainable deviance (Fig 6a) where a test model (e.g. GLM_world_) is considered in the context of GLMs that have no variable predictors of neural activity (a constant model) or use sound angle in both coordinate frames as predictors (a full model). This normalization step is critical in comparing model fit across units as deviance values alone are unbounded. In contrast, model fit is limited from 0 (indicating the sound angle provides little information about the neuron’s response) to 100% (indicating the sound angle in one coordinate frame accounts for the neuron’s response as well as sound angles in both frames). Using such an analysis, we predicted that egocentric units would have a high percentage of the full model fit by sound angles relative to the head, and a low model fit for sound angles relative to the world, and that allocentric units would show the reciprocal relationship. To test these predictions we generated a *model preference* score; the model fit for sound angles relative to the head minus the model fit for sound angles in the world (Fig 6c). Accordingly, negative values of model preference should identify allocentric units while positive values should indicate egocentric units. Neurons in which both sound angles relative to the world and head provide high model fit values may represent sounds in intermediary or mixed coordinate frames and would have model preference scores close to zero.

**Fig 6.**
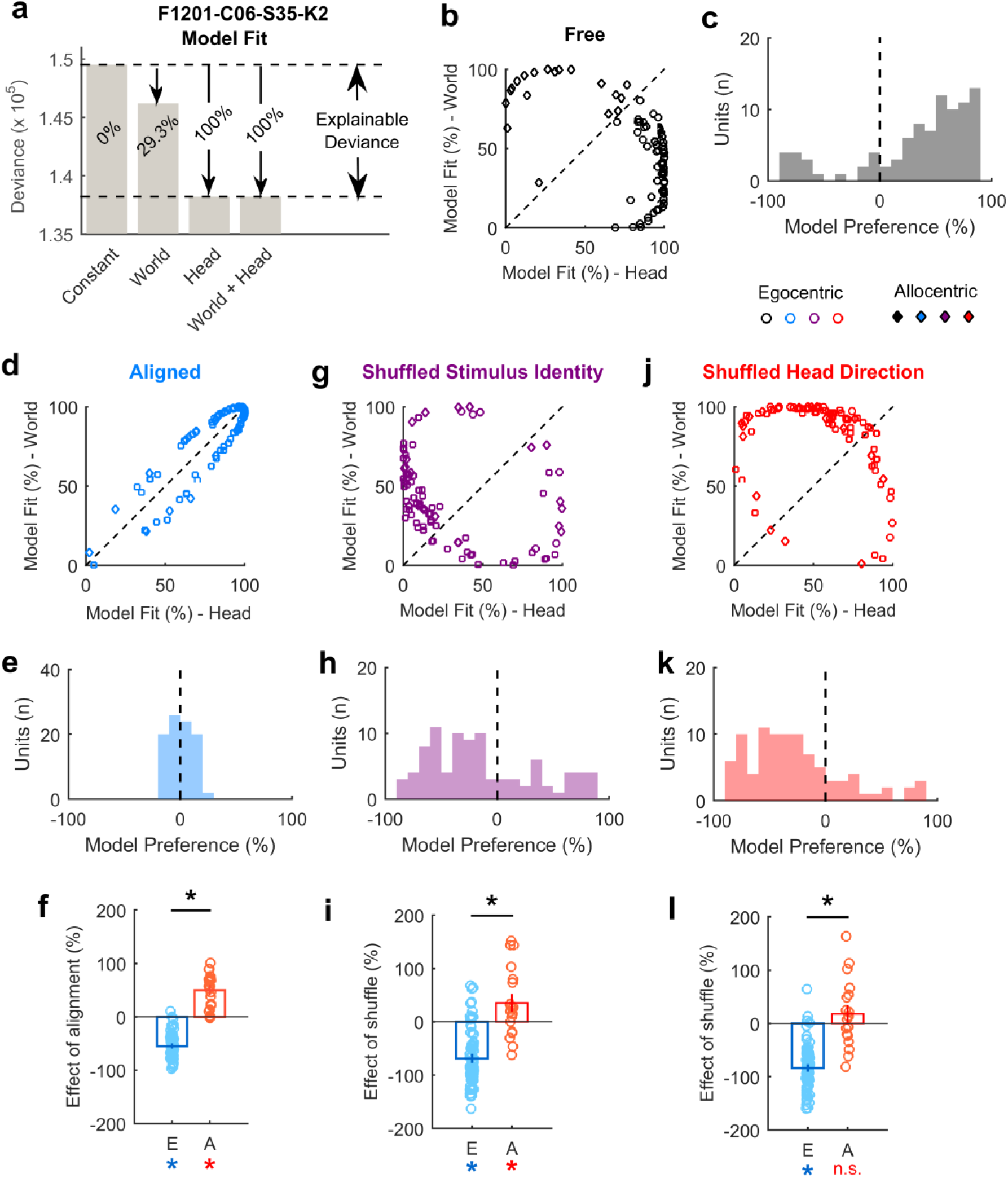
General linear modelling of spatial sensitivity. **a**, Calculation of model fit for sound angle relative to the head or world. Raw deviance values are normalized as a proportion of explainable deviance; the change in deviance between a constant and a full model. **b**, Model fit comparisons for all units when the animal was free to move through the arena. **c**, Model preference that indicates the distribution of units across the diagonal line of equality in (**b**). **d-e,** Model fit and model preference for data when the head and world coordinate frames were aligned. **f,** Change in model preference between freely moving and aligned states for egocentric (E) and allocentric (A) units. **g-h,** Model fit and model preference for freely moving data when speaker identity was shuffled. **i**, Change in model preference between unshuffled and shuffled data. **j-k,** Model fit and model preference for freely moving data when the animal’s head direction was shuffled. **i**, Change in model preference between unshuffled and shuffled data. Asterisks indicate significant differences between egocentric and allocentric subpopulations (black; p < 0.001) or significant differences from zero for a specific subpopulation (red/blue; p < 0.05).

In the space defined by model fit for sound angles relative to the head and world (Fig 6b), units clustered in opposite areas supporting the existence of distinct egocentric and allocentric neural subpopulations. This clustering was also evident in the model preference scores, which showed a distinct bimodal distribution (Fig 6c). Repeating this analysis on data in which the head and world coordinate frames were aligned (due to the animals position at the center of the speaker ring) demonstrated that model fit values for head and world models became equivalent and model preference scores were centered around zero (Fig 6d-e). Indeed, expanding our analysis from the aligned to freely varying data significantly shifted model preference for both egocentric (Fig 8f; t_7_i = 19.4, *p* = 4.02 x 10^-30^) and allocentric units (t_18_ = 6.99, *p* = 1.57 x 10^-6^), and the direction of the effect differed significantly between egocentric and allocentric units (t_89_ = 15.9, *p* = 9.71 x 10^-28^).

We performed two additional control analyses on the freely moving dataset: firstly, we randomly shuffled the speaker identity while maintaining the same information about the animal’s head direction. Randomising the speaker identity should affect the ability to model both egocentric and allocentric neurons and we would therefore predict that model fits would be worse and model preference scores would tend to zero. (I.e. shuffling would shift model preference scores in the negative direction for egocentric units and the positive direction for allocentric units). As predicted, shuffling speaker identity eliminated clustering of egocentric and allocentric units in the space defined by model fit (Fig 6g) and lead to opposing effects on model preference (Fig. 6h-i): For egocentric units, the model preference values were shifted downwards toward zero (t-test vs no change in preference: *t*_71_ = −11.0, *p* = 4.85 x 10^-17^) whereas preference values were shifted upwards towards zero for allocentric units (*t*_18_ = 2.41, *p* = 0.027). The effects of shuffling also differed between the two populations (t_89_ = −7.30, *p* = 1.17 x 10^-10^).

Secondly, we shuffled the head direction while maintaining information about speaker identity. This should cause model fit values to decline for egocentric units and should result in egocentric, but not allocentric units, shifting their model preference scores towards zero. This was indeed the case (Fig 6j-l) as shuffling head direction significantly reduced model preference for egocentric units (t-test vs zero change in model preference: *t*_71_ = −17.0, *p* = 9.65 x 10^-27^) but not allocentric units (*t*_18_ = 1.26, *p* = 0.223). The effects of shuffle by head direction also differed significantly between egocentric and allocentric units (unpaired t-test, t_89_ = −8.45, *p* = 5.09 x 10^-13^) further supporting the existence of these as distinct neural populations.

### Egocentric and allocentric units – population characteristics

For egocentric units that encoded sound location in the head coordinate frame, it was possible to characterize the full extent of tuning curves in 360° around (Fig 5a-b) despite our speaker array only spanning 180°. This was possible because the animal’s head movements were continuous across 360° and thus removed the constraints on measurement of spatial tuning imposed by the range of speaker angles used. Indeed, we were able to extend our approach further to characterize super-resolution tuning functions with an angular resolution greater than the interval between speakers (Fig 7a-b and Supplementary Fig 7). Together these findings show it is possible to deconvolve an animal’s movement from spiking activity to recover the spatial tuning of individual units with greater detail than would be possible if the subject was static. Egocentric units shared spatial receptive field properties typical of previous studies [15, 16, 18, 23]: units predominantly responded most strongly to contralateral space (Fig 7c) with broad tuning width (Fig 7e-f) that typifies auditory cortical neurons. We also found similar, if slightly weaker spatial modulation when calculating modulation depth according to Ref. [18] (Fig 7d). Across the population of allocentric units, we observed a similar contralateral tuning bias in preferred location (Fig 8a) to egocentric units and that allocentric units had relatively low modulation depths (Fig 8b). The broad modulation may not be surprising: an allocentric receptive field could presumably fall anywhere within or beyond the arena, and might therefore be poorly sampled by circular speaker arrangements. If the tuning curves measured here were in fact sampling a more complex receptive field that related to a world centered coordinate frame (Fig 1b) we would predict that the receptive fields would be correspondingly noisier. Allocentric and egocentric units were recorded at similar cortical depths (Fig 8c) and on the same cortical penetrations as both unit types were observed on 9/13 electrodes (69.2%) on which we recorded allocentric units (Fig 8d).

**Fig 7.**
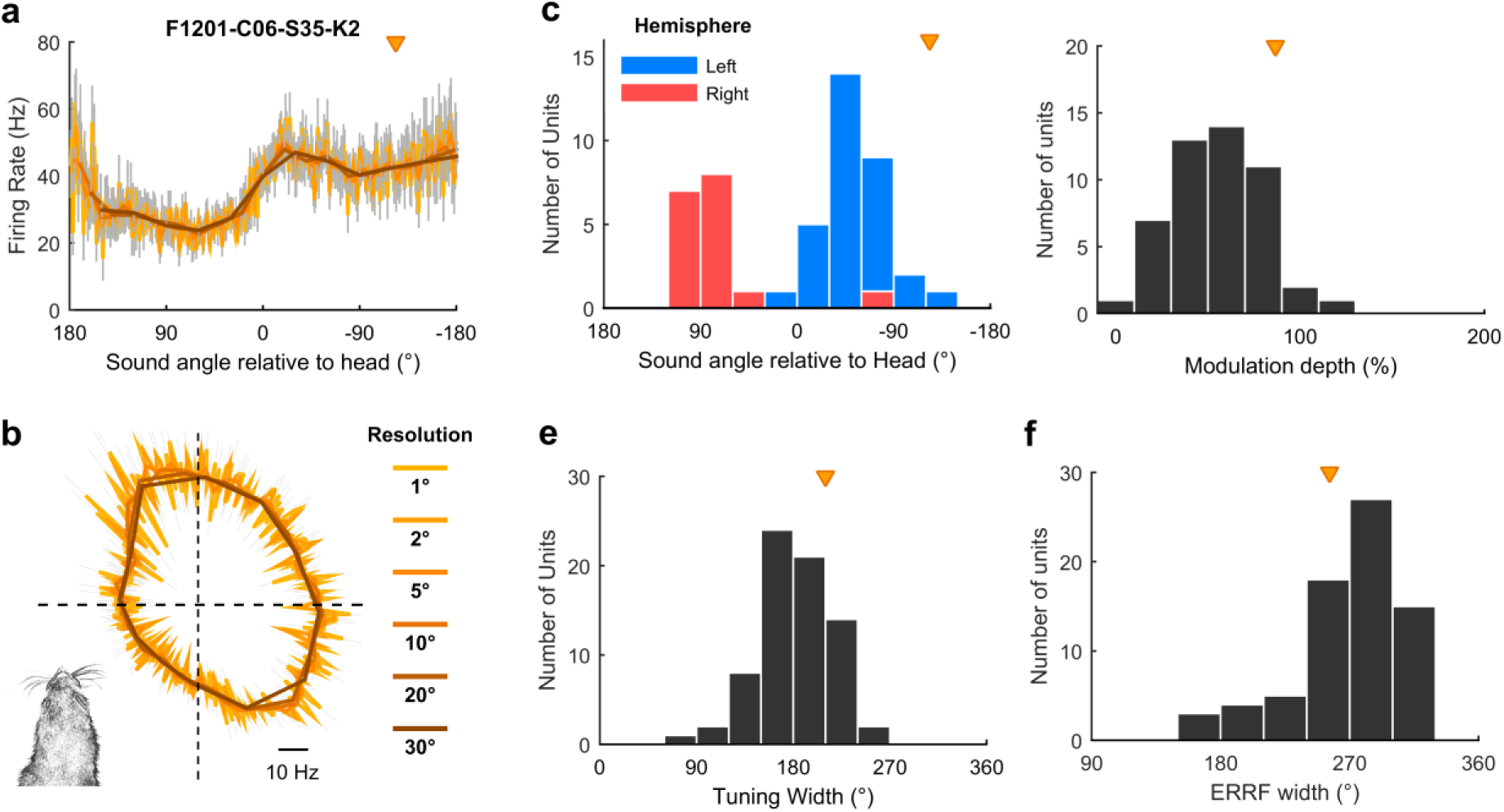
Egocentric unit characteristics. **a-b**, Spatial tuning of example egocentric unit at multiple angular resolutions. Data shown as mean ± s.e.m. firing rate plotted in Cartesian (a) or polar (b) coordinates. Triangle indicates preferred location of unit. Inset (b) illustrates the corresponding head direction onto which spatial tuning can be superimposed. **c**, Preferred location of all egocentric units (n = 72) in left and right auditory cortex. **d**, Modulation depth calculated according to [18] across 360° for units in both hemispheres. **e-f**, Tuning width (e) and equivalent rectangular receptive field width (ERRF)(f) for all units. Triangle indicates the preferred location, modulation depth, tuning width and ERRF of the example unit in (a).

**Fig 8.**
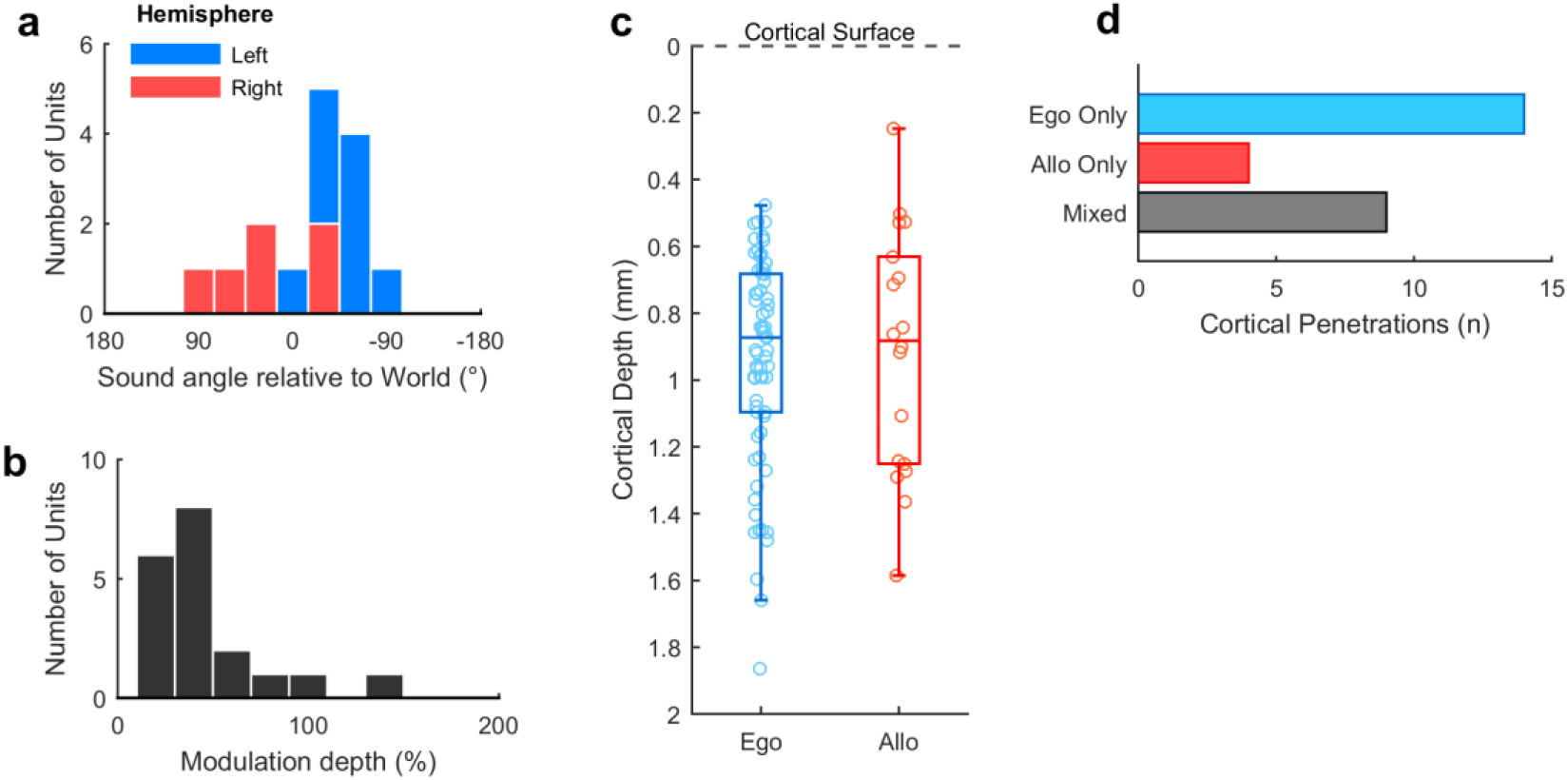
Allocentric unit characteristics. **a**, Preferred location of all allocentric units (n = 19) in left and right auditory cortex. **b**, Modulation depth calculated across 180° for units in both hemispheres. **c,** Comparison of cortical depth at which egocentric and allocentric units were recorded. Ferret auditory cortex varies in thickness between 1.5 and 2 mm and electrode depths were confirmed histologically (Supplementary Fig. 2). **d**, Number of cortical penetrations on which we recorded only egocentric units, only allocentric units or a combination of both (mixed). All 92 spatially tuned units were recorded on 27 unique electrodes, with most units at a single cortical location recorded at different depths as electrodes descended.

### Timing of spatial information

Our findings suggested the existence of distinct egocentric and allocentric populations of spatially tuned units. As these subpopulations were functionally defined, we hypothesized that differences between allocentric and egocentric units should only arise after stimulus presentation. To test this, we analyzed the time course of unit activity in a moving 20 ms window and compared model fit and model preference of egocentric and allocentric units (defined based on the AIC analysis above) using cluster-based statistics to assess statistical significance [24]. Model fit for sound angles relative to the head was greater for egocentric than allocentric units only between 5 and 44 ms after stimulus onset (Fig 9a, *p* = 0.001996). Model fit for sound angles in the world was greater for allocentric than egocentric units only between 6 and 34 ms after stimulus (Fig 9b, *p* = 0.001996). Model preference of the two populations diverged only in the window between 4 and 43 ms after stimulus onset (Fig 9c, *p* = 0.001996). We observed no differences between sub-populations before stimulus onset or when coordinate frames were aligned (Supplementary Figure 8). Thus the distinction between egocentric and allocentric units reflected a stimulus-evoked functional division that was only observed during free movement.

**Fig 9.**
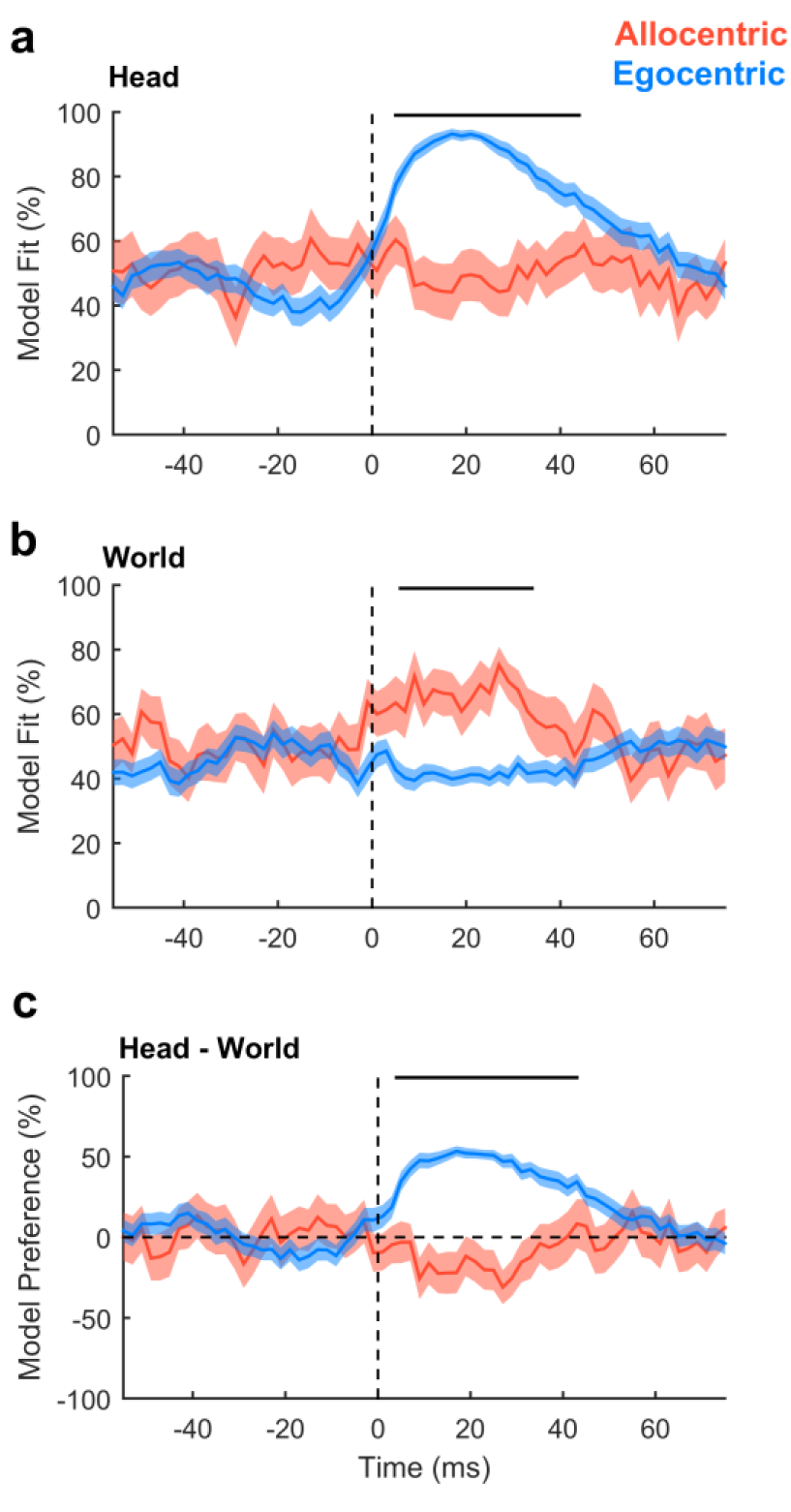
Egocentric and allocentric population distinctions in time. **a**, Model fit for predicting neural activity from sound angles relative to the head. **b**, Model fit for predicting neural activity from sound angles in the world. **c**, Model preference. Data shown as mean ± s.e.m. for egocentric and allocentric unit populations. Black lines indicate periods of statistical significance (cluster based unpaired t-test, *p* < 0.05).

### Population representations of space

We next asked how auditory cortex neurons behaved as a population when spatial tuning was compared across head directions. In contrast to individual units, population activity more closely reflects the large-scale signals observed in human studies using electroencephalography (EEG) to distinguish coordinate frame representations [11, 12]. As would be expected from the predominance of egocentric units, we found that tuning curves for both left and right auditory cortical populations (n = 64 and 28 units respectively) were consistent within the head but not world coordinate frame (Fig 10). Thus the allocentric units we find here are sufficiently rare as to be masked in overall population readouts of spatial tuning, potentially accounting for conflicting findings of coordinate frame representations from EEG recordings.

**Fig 10.**
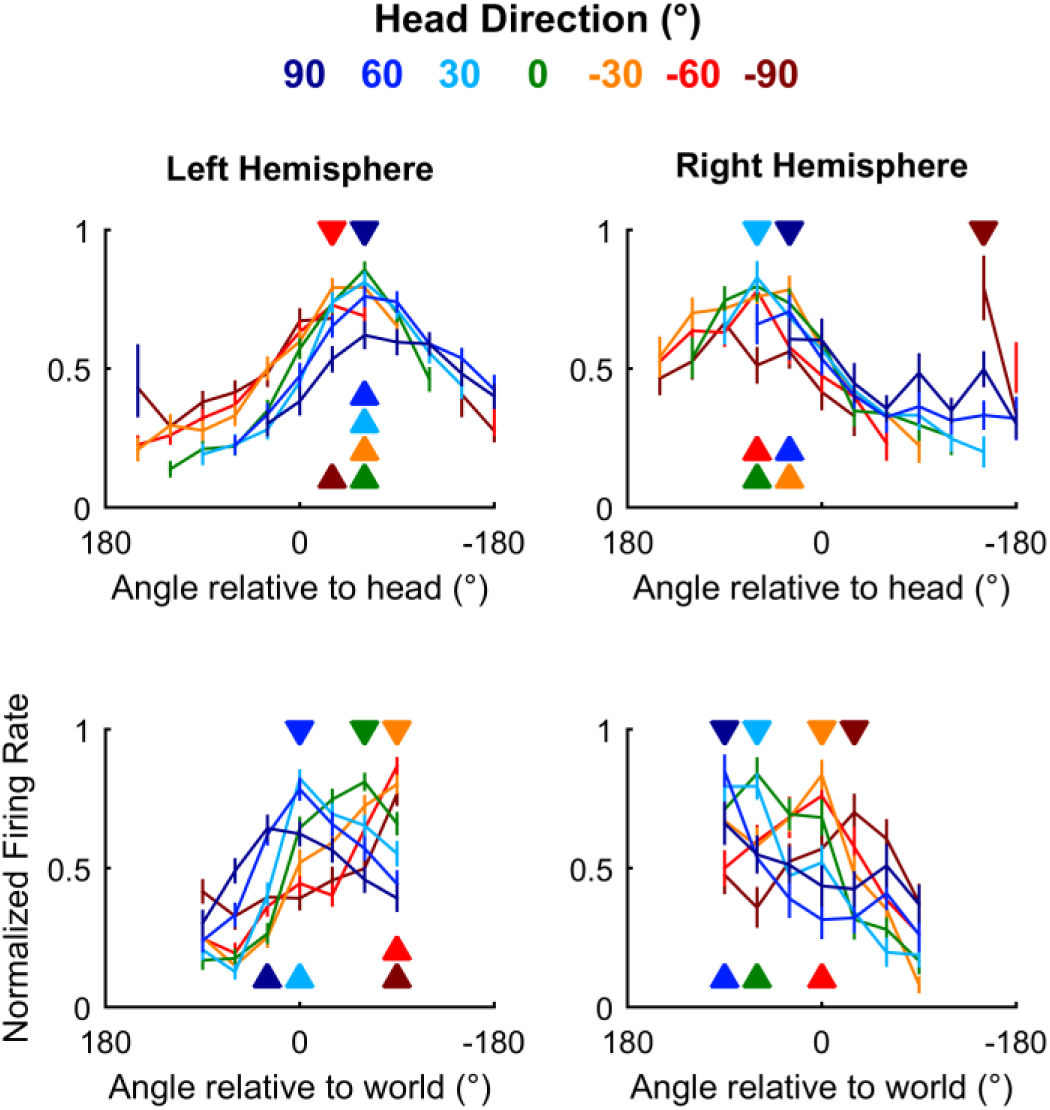
Auditory cortical tuning. Population tuning curves plotted across head direction for mean (± s.e.m.) normalized response of all units in left and right auditory cortex; filled triangles indicate sound angle of maximum response at each head direction.

### Distance modulation of cortical neurons and spatial tuning

Studying auditory processing in moving subjects allowed us to resolve coordinate frame ambiguity so that we could identify the spaces in which neurons represent sound location and define populations of egocentric and allocentric units. However recording in freely moving subjects also made it possible to go beyond angular measurements of the source location and address how neurons represented the distance of sound sources. Though often overlooked, distance is a critical component of egocentric models of neural tuning as absolute sound level and the acoustic cues supporting head-centered localization vary with distance between near-field (< 1 meter) and far-field conditions [25, 26]. In our study, the distance between head and sound source ranged from 10 cm (the minimum distance imposed by the arena walls) to 40 centimetres, with only 3.37% of stimuli (mean across 57 test sessions) presented at greater distances (Fig. 2g) and thus all stimuli were presented in the near field. Clicks were presented from each sound source with a roving level across 6 dB and the sound level at the head ranged from 48 to 67 dB SPL (Fig. 2h; median minimum and maximum sound levels across sessions).

Spatial tuning was observed at all distances studied in both egocentric (Fig 11a) and allocentric units (Fig 11b) however modulation depth increased with distance for egocentric units (ANOVA, F_2_,_216_ = 3.45, p = 0.0334). Pairwise post-hoc comparisons showed that modulation depth was largest for sounds at greatest distances (Fig 11c) though only significantly so when comparing modulation depth measured for sound presented 20 to 30 cm from the animal with sounds presented 30 to 40 cm away (*t_72_* = −3.54, *p* = 0.0279). In contrast, modulation depth did not change significantly with distance for allocentric units (Fig 11d, F_2_, _57_ = 0.962, p > 0.1).

**Fig 11.**
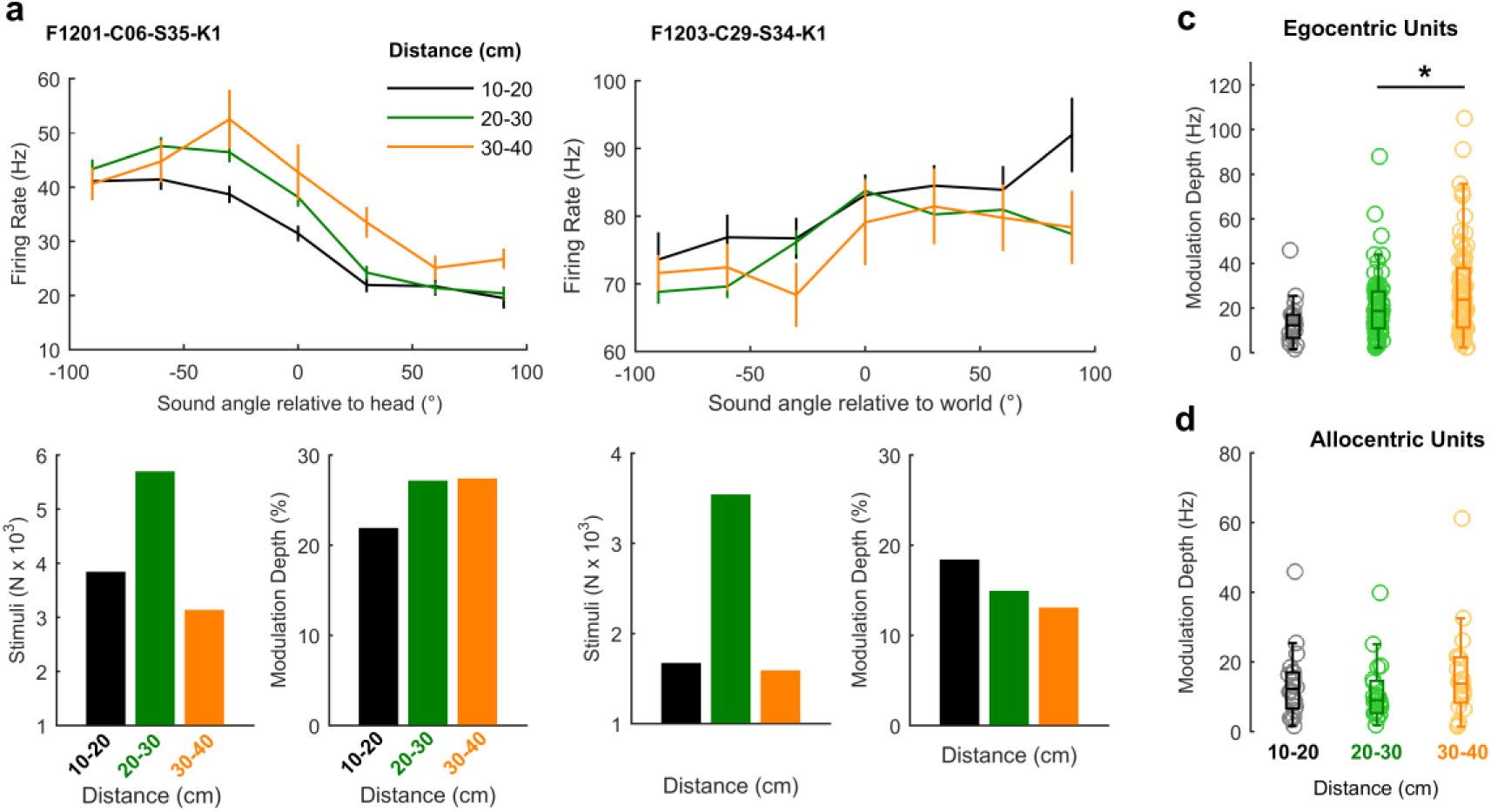
Spatial tuning across distance. **a-b**, Tuning curves of an egocentric (**a**) and allocentric (**b**) unit obtained with sound sources at varying distances from the animal’s head. Bar plots show the number of stimuli and modulation depth for each tuning curve. **c-d** Distributions of modulation depth measured across distance for egocentric and allocentric sub-populations. Asterisk indicates significant pair-wise comparison (p < 0.05).

### Speed modulation of cortical neurons and spatial tuning

Studying auditory processing in moving subjects also allowed us to investigate how speed of head movement (Fig. 2f) affected neural activity. Movement is known to affect auditory processing in rodents [27-29] but its effects on spatial representations of sound location and also on auditory cortical processing in other phyla such as carnivores remain unknown.

To address auditory cortical processing first, we asked how many of our recorded units (regardless of auditory responsiveness or spatial modulation) showed baseline activity that varied with speed. For each unit, we took all periods of exploration (excluding the 50 ms after each click onset) and calculated the speed of the animal at the time of each action potential. We then discretized the distribution of spike-triggered speeds to obtain spike counts as a function of speed and normalized spike counts by the duration over which each speed range was measured. This process yielded a speed-rate function for baseline activity (Fig 12a). We then fitted an exponential regression curve to each function (Fig 12b) and plotted the correlation (R^2^) and regression coefficients (β) of each curve to map the magnitude and direction of association between speed and baseline activity (Fig 12c). Across the recorded population, we saw both positive and negative correlations representing units for which firing rate increased or decreased respectively with speed. However a significantly larger proportion of the population increased firing rate with speed across all units (t-test vs. 0; *t*_308_ = 3.77, *p* = 1.97 x 10^-4^). This was also true if we considered only sound responsive units (t_267_ = 5.15, *p* = 5.17 x 10^-7^) or only spatially tuned units (*t*_91_ = 4.12, *p* = 8.41 x 10^-5^).

**Fig 12.**
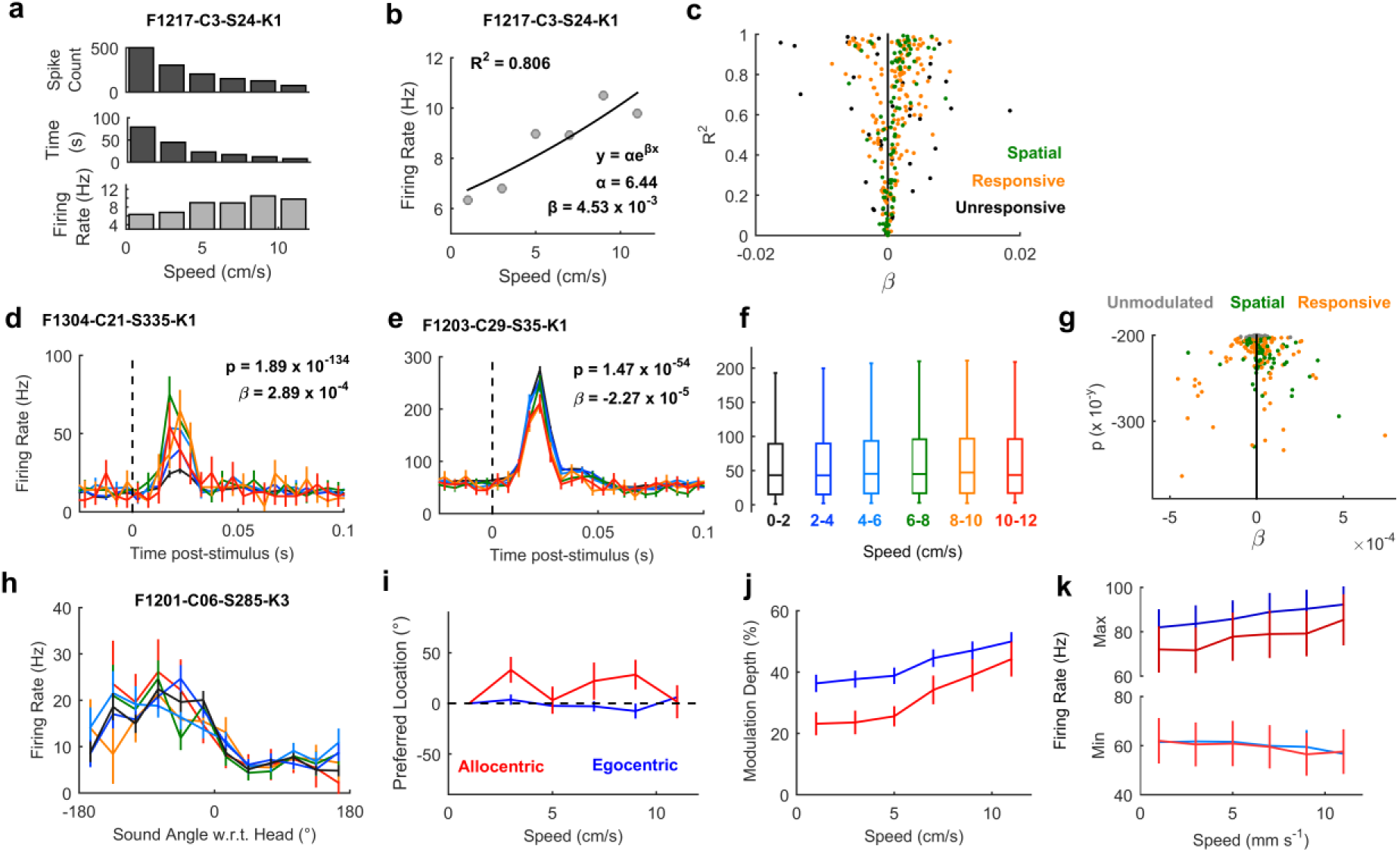
Speed related auditory cortical activity and sensory processing. **a** Example calculation of speed-related modulation of baseline firing of one unit using reverse correlation. **b**, Example speed-firing rate function summarized using regression (β) and correlation (R^2^) coefficients for the same unit as (**a**). **c**, Population distribution of regression and correlation coefficients showing the predominance of units with increasing speed-rate functions (β >0). **d-e**, Peristimulus time histogram of sound evoked responses for units that were enhanced (**d**) or suppressed (**e**) at increasing speeds. **f,** Box plots showing distributions (median and inter-quartile range) of evoked firing rates in response to clicks across speed for all sound responsive units. **g**, Population distribution of regression coefficients (β) and model fit (analysis of deviance p values) for all sound-responsive units. Units for which speed was not a significant predictor of neural activity (p > 0.001) denoted in grey. **h**, Spatial tuning curve for one unit for clicks presented at different head movement speeds. **i-k**, Change in preferred location (**i**), modulation depth (**j**) and min / max firing rates (**k**) of egocentric and allocentric units as a function of speed. Data for d-e and h-k shown as mean ± s.e.m.

We also observed both speed-related suppression (Fig. 12d) and enhancement (Fig 12e) of sound-evoked responses in individual units. For each individual unit, we characterized the relationship between head speed and single trial firing rates (averaged over the 50 ms post-stimulus onset) using a general linear model that measured both the strength (analysis of deviance *p* value) and direction (model coefficient, β) of association. Positive β values indicated an increase in firing rate with increasing speed whereas negative β values indicated a fall in firing rate with increasing speed. Thus the direction of the relationship between firing rate and speed was summarized by the model coefficient, allowing us to map the effects of movement speed across the population (Fig.12f). For 199/268 sound-responsive units (74.3%), speed was a significant predictor of firing rate (analysis of deviance vs. a constant model, *p* < 0.001) however the mean coefficient value for movement sensitive units did not differ significantly from zero (t_199_ = 0.643, *p* = 0.521). This suggests the sound-responsive population was evenly split between units that increased or decreased firing with speed. We noted that a significantly greater proportion of spatially modulated units (74/92, 80.4%) had sound evoked responses that were sensitive to speed than units that were either not spatially modulated or for which we had insufficient sample sizes to test spatial modulation (125/176, 71.0%)(Chi-squared test, *χ^2^* = 3.96, *p* < 0.05). For those 74 spatially modulated and speed sensitive units, coefficients were mostly larger than zero (mean ± s.e.m. = 2.73 x 10^-5^ ± 1.53 x 10^-5^) however this effect was not statistically significant (*t*_73_ = 1.80, *p* = 0.076). For the remaining speed sensitive units, the mean coefficients was closer to zero (mean ± s.e.m. = −5.02 x 10^-6^ ± 1.49 x 10^-5^).

Lastly, we asked if speed affected spatial tuning. Spatial tuning could be observed at all speeds of movement in spatially tuned units (Fig. 12h) and the preferred locations of units did not vary systematically with speed (Fig 12i). For neither egocentric nor allocentric units was there a significant effect of speed in an ANOVA on preferred locations (egocentric: *F*_5_, _432_ = 0.53, *p* = 0.753; allocentric: F_5_, i_08_ = 1.53, p = 0.188). However we did observe significant changes in modulation depth (Fig 12j) both for egocentric (F_5_, _432_ = 4.91, *p* = 0.0002) and allocentric units (F_5_, _108_ = 5.09, *p* =0.0003), indicating that spatial tuning was sharpest when the head was moving fastest. Change in modulation depth resulted from both a gradual suppression in minimum and enhancement in maximum firing rates with speed (Fig 12k). However none of these changes were significant in comparisons across speed (ANOVA with speed as main factor, p > 0.5) indicating that it was only through the aggregative change in responses to both tuned and untuned locations that modulation depth increased with speed.

## Discussion

Here we have shown that by measuring spatial tuning curves in freely moving animals it is possible to unambiguously demonstrate the coordinate frames in which neurons define space and thus sound location. For the majority of auditory cortical neurons, we found egocentric tuning that confirm the broadly held but untested assumption that sound locations are represented (for the most part) in head-centered coordinates. We also identified a novel sub-population of units with allocentric tuning, whose responses were spatially locked to sound location in the world across movement. Such allocentric units were consistently identified and functionally distinct from egocentric units across multiple analyses, and were observed in multiple subjects, suggesting that multiple coordinate frames are represented at the auditory cortical level. Finally we used the advantages of a freely moving subject to explore the dependence of neural activity and spatial tuning on sound source distance and the speed of head movements, demonstrating significant roles for both that emphasise the importance of studying active sensing in unrestricted animals.

Our findings reveal that auditory cortex represents sound location in at least two coordinate frames during free movement. Our identification of allocentric representations in which sound location is consistently reported in the world across self-motion supports the role of auditory cortex in parsing an auditory scene into distinct objects [30, 31]. When we turn our heads our perception of a sound source remains stable, despite the fact that in physical terms it actually moves relative to us. Stable perception requires that changes in sensory input that result from self-generated motion are disambiguated from those that result from source motion [32]. The allocentric representations described here fulfil this role and thus offer a potential correlate of a stable perceptual world at the neural level.

A key question following on from the discovery of allocentric units is where they reside in cortex. The units recorded here were located on the same electrodes and cortical depths as egocentric units, in the primary and posterior regions of auditory cortex. However, targeting the low-frequency reversal where primary and posterior fields meet and the use of electrode arrays as chronic behavioral implants prevented us from mapping the precise tonotopic boundaries necessary to attribute units to specific cortical subfields [33]. Future use of denser sampling arrays may enable cortical mapping in behaving animals and thus precise localization of allocentric units. We targeted primary and posterior auditory cortex as these areas are likely to be sensitive to inter-aural timing cues, and the animals involved in this work were also trained to discriminate non-spatial features of low frequency sounds in another study (Town et al, Unpublished). Posterior regions may correspond to part of the ‘what’ pathway in auditory processing whereas the anterior ectosylvian gyrus may correspond to the ‘where’ pathway in which spatial tuning is more extensive [34, 35]. It is thus likely that the coordinate frames represented in our population (where only 49% of units were spatially sensitive) may be more ubiquitous in anterior regions of ferret auditory cortex. Indeed given sensorimotor and cross-modal coordinate frame transformations are a key feature of activity in parietal cortex [10], it is possible that allocentric representations exist beyond auditory cortex.

While the presence of allocentric receptive fields in tonotopic auditory cortex is surprising, the existence of allocentric representations has been predicted by behavioural studies in humans [6, 13]. Furthermore, coordinate transformations occur elsewhere in the auditory system [36, 37] and behavioral movements can influence auditory subcortical and cortical processing [27, 28]. Perhaps most importantly, vestibular signals are integrated into auditory processing already at the level of the cochlea nucleus [38] allowing the distinction between self and source motion [22]. Auditory-vestibular integration, together with visual, proprioceptive and motor corollary discharge systems, provides a mechanism through which changes in head direction can partially offset changes in acoustic input during movement to create allocentric representations. However an outstanding question is how head position is also integrated into auditory processing. The navigation system within the medial temporal lobe (or its carnivore equivalent) is a leading candidate given the connections between entorhinal and auditory cortex [39, 40]; however the functional interactions between these areas and their potential contributions to allocentric processing remain to be addressed.

In addition to identifying allocentric units, we also show that an animal’s movement can be successfully deconvolved from auditory responses to measure head-centered egocentric tuning during behavior. While we used speakers placed at 30° intervals across a range of 180°, we were nonetheless able to characterize spatial tuning of egocentric units around the full circumference of the head (i.e. Fig. 3c). This illustrates the practical benefit of using freely moving subjects to characterize head-centered spatial tuning as the animal’s movements generate the required variation in sound angle relative to the head to map spatial tuning and thus the number of sound sources needed was reduced. Furthermore, as the animal’s head direction was continuous, the stimulus angle was also continuous and thus it was possible to measure spatial tuning at higher resolutions than would otherwise be possible with speakers at set intervals. In contrast to egocentric tuning, our ability to map allocentric receptive fields was limited by the speaker arrangement that only sparsely sampled world coordinates (Fig 2a). This may in part explain the low number of allocentric units in our population and denser sampling of the world may reveal unseen allocentric tuning – for example in the 94 units we recorded that were not spatially modulated by sound locations in the world that we tested. While a full 360° speaker ring may offer a minor improvement in sampling density, the radial organization of the ring remains a suboptimal design for sampling rectangular or irregular environments. To fully explore the shape of allocentric receptive fields will require denser, uniform speaker grids or lattices in environments through which animals can move between sound sources.

A notable property of egocentric units was the relationship between modulation depth of spatial tuning and distance, which was also absent in allocentric units. Here this distance sensitivity may largely be driven by sound level at the head, although other auditory and non-auditory cues can affect distance perception [41-45]. The lower modulation depth at closer distances may result from disruption to acoustic cues such as inter-aural level differences to which cortical neurons are sensitive [46] but that become more complex in the near field (distances of less than one meter) [25, 26]. However while acoustic cues in the near field are complex, cues such as inter-aural time differences remain informative about sound location [25, 47] and could support sound localization in the near field in both ferrets and humans [48](Wood et al., unpublished). It may not be surprising therefore that we were able to record spatial tuning in large numbers of units despite presenting all sounds from sources within the near field. In future it will be critical to validate our findings for sound sources at greater distances in the far field where sound localization has been more widely studied.

The use of moving subjects also allowed us to study the effects of movement speed on auditory processing and spatial tuning. In contrast to other studies in freely behaving animals that reported movement-related suppression of activity [27-29] we found that neurons tended to fire more strongly when the animal moved faster (Fig 12). One reason for this difference may lie in the behavior measured: Other investigators have covered a diverse range of actions including locomotion in which both the head and body move and self-generated sounds are more likely. We only considered the head speed of an animal and did not track the body position that would allow distinction of head movements from locomotion (which was relatively limited given the size of the animal and the arena). It is thus likely that much of the variation in speed we observe is a product of head movement during foraging without locomotion and thus with relatively little self-generated sound. The behavior of our subjects may therefore present distinct requirements for auditory-motor integration that result in distinct neural effects.

We also observed that spatial modulation was also greater when the animal was moving faster, which may be consistent with the sharpening of tuning curves during behavioral engagement [15]. While sharpening of engagement-related spatial tuning was linked to a reduction in spiking responses at untuned locations, we observed non-significant decreases in peak and minimum firing rates suggesting that the mechanisms underlying speed-related modulation of spatial tuning may be subtly different. At the acoustic level, faster movements provide larger dynamic cues [2, 3] that improve sound localization abilities in humans [3, 49, 50] and may explain the increase in modulation depth of units at greater speeds observed here.

In summary, we recorded spatial tuning curves in freely moving animals to resolve coordinate frame ambiguity and demonstrated distinct populations of egocentric and allocentric units. In addition to revealing the existence of allocentric representations, self-generated movement offers advantages for extending the study of egocentric sound processing and the interaction between spatial representations and motor activity. Together this shows the value of studying neural processing in the freely moving state in which most animals naturally behave.

## Methods

### Simulated spatial receptive fields

#### Egocentric

Egocentric tuning described the relationship between spike probability (P) and sound source angle relative to the midline of the subject’s head (*θ_HS_*) and was simulated in Matlab (MathWorks) using a Gaussian function:

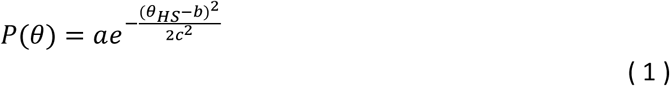

In the example shown in figure 1a, parameters (a = 1.044, b = 0° and c = 75.7°) were determined by manual fitting to find values for which egocentric and allocentric tuning matched qualitatively. The theta domain was between ±180° binned at 1° intervals and distance of sound sources was not included in the simulation.

#### Allocentric

Allocentric tuning describes the relationship between neural output (reported here as spike probability; *P*) and sound source position within the world measured in Cartesian (*x,y*) coordinates. Spatial tuning was simulated as the dot product of spike probability vectors returned from functions defined separately for positions on the x- and y-axes:

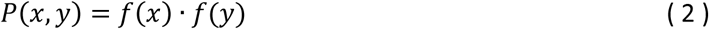

In figure 1b, logistic probability functions were used for both dimensions:

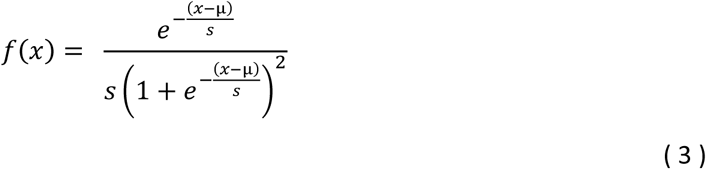

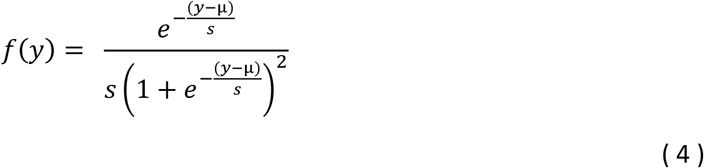

With μ = 1000 mm and s = 400 mm for the x-axis, and μ = 0 mm and s = 1000 mm for the y-axis. For both axes, spike probability vectors were generated for domains between ± 1500 mm binned at 2 mm intervals.

#### Stimulus presentation, head pose and movement

The position and orientation of the subject’s head within the world was described as a coordinate frame transform composed of a translation vector between origins (indicating the head position) and a rotation matrix between axes (indicating the head direction). Stimuli were presented on each time step of the simulation from each speaker in a ring at 10° intervals, 1000 mm from the origin of the world coordinate frame. As the simulation was deterministic, each stimulus was presented to static simulations only once to calculate the response. When simulating motion, a ‘pirouette' trajectory was constructed in which the subject’s head translated on a circular trajectory (radius = 50 mm; angular speed = 30° per time-step) while simultaneously rotating (angular speed = 10° per time-step) for 7200 stimulus presentations (Supplementary Video 2). For each stimulus presentation, the stimulus angle was calculated relative to both the midline of the head and the vertical axis of the arena (Supplementary Fig 9). Simulation responses were quantized in 1° bins.

### Animals

Subjects were five pigmented female ferrets (1-5 years old) trained in a variety of psychophysical tasks that did not involve the stimuli presented or the experimental chamber used in the current study. Each ferret was chronically implanted with Warp-16 microdrives (Neuralynx, MT) housing sixteen independently moveable tungsten microelectrodes (WPI Inc., FL) positioned over middle and posterior fields of left or right auditory cortex. Details of the surgical procedures for implantation and confirmation of electrode position are described elsewhere[51].

Subjects were water-restricted prior to testing, during which they explored the experimental arena to find freely available sources of water. On each day of testing, subjects received a minimum of 60ml/kg of water either during testing or supplemented as a wet mash made from water and ground high-protein pellets. Subjects were tested in morning and afternoon sessions on each day for up to three days in a week (i.e. a maximum of six consecutive testing sessions); on the remaining weekdays subjects obtained water in performance of other psychophysical tasks. Test sessions lasted between 10 and 40 minutes and were ended when the animal lost interest in exploring the arena.

The weight and water consumption of all animals was measured throughout the experiment. Regular otoscopic examinations were made to ensure the cleanliness and health of ferrets’ ears. All experimental procedures were approved by a local ethical review committee and performed under license from the UK Home Office and in accordance with the Animals (Scientific Procedures) Act 1986.

### Experimental Design and Stimuli

In each test session, a ferret was placed within a D-shaped arena (Fig. 2a, rectangular section: 35 x 30 cm [width x length]; semi-circular section: 17.5 cm radius; 50 cm tall) with seven speakers positioned at 30° intervals, 26 cm away from a central pillar from which the animal could find water. The periphery of the circular half of the arena was also fitted with spouts from which water could be obtained. Animals were encouraged either to explore the arena by delivery of water at all spouts, or to hold their head at the center spout by restricted water delivery at this location. The arena and speakers were housed within a sound-attenuating chamber lined with 45 mm sound-absorbing foam.

During exploration (n = 57 test sessions), clicks were presented from each speaker with random inter-stimulus intervals (250 - 500 ms). The instantaneous energy of clicks minimized dynamic cues, simplifying neural analysis and comparisons with other work on spatial encoding. Clicks were presented at 60 dB SPL when measured from the center of the arena using a measuring amplifier (Bruel & Kjaer 2636). However because sound level varied across the arena, we roved sound levels over a ±6 dB range to counter changes in level arising from differences in position of the head within the sound field. The frequency response of each speaker (Visaton SC 5.9) was measured using golay codes [52] and compensated for to produce a flat spectral output between 20 Hz and 20 kHz. Stimulus location and water delivery were independent and subjects were not required to attend to stimuli in order to find water rewards. To avoid characterizing neural responses to the sound of solenoid control signals, stimulus presentation and water reward were delivered in separate alternating time windows; water was delivered in a short period of 1 to 2 seconds when each solenoid was rapidly opened (100 ms duration) with a 10 second interval between delivery windows in which click stimuli were presented. Sessions typically lasted approximately 15 - 20 minutes (median = 16.5 minutes; range = 6.15 - 48.0 minutes) in which several thousand stimuli could be presented (median = 1984; range = 304 - 3937).

### Head Tracking

During exploration of the experimental arena, the animal’s head position and orientation were tracked using two LEDs (red and green) placed along the midline of the head and recorded using an RV2 video acquisition system (TDT) sampling at 30 frames per second and synchronized with the electrophysiology recording hardware. For each video frame, the red and green LED positions were identified in Matlab from a weighted image in which the channel color of the target LED was positively weighted and all other channels negatively weighted. Each LED position was then taken as the center of the largest cluster of pixels containing the maximum weighted value. To maximize the frame rate of the camera, we recorded with a low exposure time (10-20 ms). Lower exposure also improved LED identification by reducing signal intensity in the background of each frame.

In cases where an LED went out of view of the camera (usually due to the roll or pitch of the head, or the recording cables obscuring the LED), the maximum weighted value identified as the LED would be a random point within the arena resulting either from a weak reflection or image noise. To remove such data, we set a minimum intensity threshold based on the distribution of maximum values in weighted images across all frames. In cases where the LED intensity failed to match the specified threshold, the LED position was noted as missing. To compensate for missing data, we estimated LED positions across runs of up to a maximum of ten frames (333 ms) using spline interpolation. Longer runs of missing data were discarded.

We then mapped each LED position in the image (M) into the behavioral arena to give the new position *N* using the transformation:

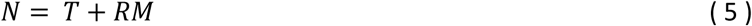

Where *T* is the translation between the origin of the image coordinate frame (i.e. pixel [0,0]) and the origin of the arena coordinate frame (the center of the arena). And, *R* is the three-dimensional rotation matrix describing the rotation between the arena and image coordinate frames. *T* was obtained by manually identifying the pixel closest to center of the arena (i.e. the equidistant point between all speakers) in a calibration image captured at the start of each test session. *R* was estimated from singular value decomposition using the position and distance between known points in the arena (also identified manually from each calibration image). Here we estimated a 3D rotation matrix to take into account the position of the camera relative to the arena (i.e. above the arena rather than below). All coordinate frames were represented using the right-hand rule (i.e. positive values for counter-clockwise rotation about the z-axis) to ensure consistency with the *atan2* function.

Within the arena, the animal’s head position 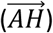 was then calculated as the mid-point between the LEDs within the arena and was used to define the origin of the head-centered coordinate frame (Supplementary Fig 9). The animal’s head direction (*θ_HA_*) was calculated from the two argument arctangent function (atan2) of the vector between LEDs that defined the midline (Y-axis) of the head-centered coordinate frame 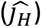. The *Z* axis was undefined by the tracking system as we only measured two points (red and green LEDs) with a single camera; this leads to ambiguity about the pitch and roll of the head. To compensate for this deficiency we assumed that when LEDs were visible, the *XY* plane of the head always matched the plane of the arena floor and that the Z-axis of the head was orthogonal to this plane and oriented towards the camera. Such assumptions are justified by the properties of the tracking system - as the head rolls or pitches away from the assumed conditions, it becomes impossible to identify both LEDs within the image due to the limited angular range of the each diode. Therefore tracking was impossible (in which case data was discarded) in the same conditions in which our assumptions became untenable.

By using the frame times recorded on the device, it was possible to create a time series of head position and direction within the arena that could be compared to the spiking pattern of neurons. We used the inter-frame interval and change in position of the head origin, smoothed with a nine-point Hann window to calculate the speed of head movement.

### Neural Recordingy

Neural activity in auditory cortex was recorded continuously throughout exploration. On each electrode, voltage traces were recorded using TDT System III hardware (RX8 and RZ2) and OpenEx software (Tucker-Davis Technologies, Alachua, FL) with a sample rate of 25 kHz. For extraction of action potentials, data were bandpass filtered between 300 and 5000 Hz and motion artefacts were removed using a decorrelation procedure applied to all voltage traces recorded from the same microdrive in a given session [53]. For each channel within the array, we identified candidate events as those with amplitudes between −2.5 and −6 times the RMS value of the voltage trace and defined waveforms of events using a 32-sample window centered on threshold crossings. Waveforms were then interpolated (128 points) and candidate events combined across sessions within a test run for spike sorting. Waveforms were sorted using MClust (A.D. Redish, University of Minnesota, http://redishlab.neuroscience.umn.edu/MClust/) so that candidate events were assigned to either single-unit, multi-unit clusters or residual hash clusters. Single units were defined as those with less than 1% of inter-spike intervals shorter than 1 millisecond. In total 331 units were recorded, including 116 single units (35.1%).

### Tracking unit identity across recording sessions

Through the experiment, electrodes were descended progressively deeper into cortex at intervals of 50 - 100 μm to ensure sampling of different neural populations. At most recording sites, we tested animals on multiple sessions (1-6 sessions) across several (1-3) consecutive days. Conducting test sessions over multiple days makes possible the recording of different units at a single recording site over time (i.e. through electrode drift, gliosis etc.). To constrain our analysis to units with a consistent identity we tracked the waveform of recorded units across sessions within a test run. Our rationale was that a unit should have a constant waveform shape across test sessions and any differences in waveform shape should be small relative to differences in the waveforms of units measured on other electrodes or at other depths by the same electrode. Thus for one test session at a given recording site, we calculated the Euclidean distance matrix between the mean waveform recorded on that session (W_Test_) and the mean waveform recorded on each additional session at the same recording site (D_Test_). We also calculated the distances between W_Test_ and the mean waveform recorded for every session at different recording sites (D_Controļ_). D_Contro_| provided null distributions for waveform distances between pairs of neurons known to have separate identities (due to the spatial separation between electrodes at recording sites [>50 μm in depth, >500 μm laterally]). For a given waveform, we then calculated the statistical probability of observing distances between test waveform and waveforms *at the same recording site* given the distribution of distances between test waveforms and waveforms *at other recording sites.* For waveforms exceeding statistical significance (t-test; *p* < 0.05, Bonferroni corrected for the number of sessions conducted at the recording site), we concluded that the same neuron or neuronal population was recorded.

For runs of test sessions, we took the longest continuous run for which waveform distances were significantly smaller than expected by chance. The majority of units tested more than once could be tracked over all sessions tested (72.4%: 126 / 174 units) although the number of neurons tracked fell off with time.

### Data Analysis

During exploration we characterized sound evoked responses from auditory cortical units. Each click stimulus and the concomitant neural response could be related to controlled variables determined by the experimental design and measured variables observed from head tracking. Controlled variables were the position of the sound source relative to the arena and sound source level in dB SPL whereas measured variables were the position and direction of the head relative to the arena, as well as head speed. Controlled and measured variables were combined to determine several experimental parameters: Stimulus position relative to the head was calculated as the vector 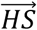:

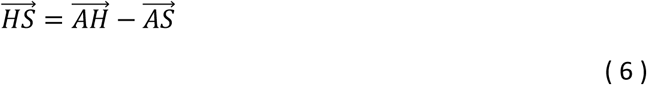

Where 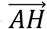 is the vector from arena origin to head origin and 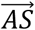 is the vector from arena origin to the sound source. Stimulus angle relative to the head (*θ_HS_*) was calculated by subtracting head direction in the arena (*θ_HA_*) from the stimulus angle relative to the origin of the head coordinate frame:

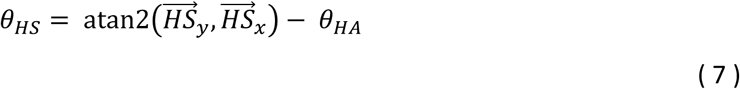

The distance between head and stimulus was calculated as the magnitude of 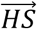 and expressed as a ratio relative to the distance between arena and stimulus to calculate sound level at the head:

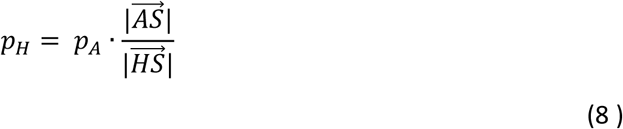

Where P_H_ and P_A_ are sound pressures at the head and center of the arena expressed in pascals and sound level is expressed in dB SPL:

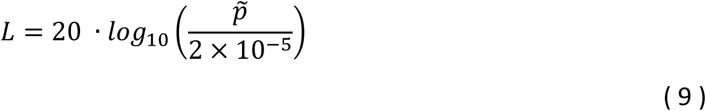

Sound level was calibrated to 60 dB SPL (0.02 Pa) at the center of the arena.

For each variable we calculated the value at the time of stimulus presentation (i.e. with a lag of 0 ms) and contrasted these values with the spiking responses of neurons. To study encoding of stimulus features (both measured and control variables) by neurons, single trial responses of individual units were summarized as the mean firing rate 0 - 50 ms after stimulus onset. This window was sufficiently long to characterize the response of units (e.g. Fig 3) but also short enough that changes in head direction and position during the analysis window were small (Supplementary Fig 3). Sound-responsive units (268/336) were first identified as those with evoked firing rates that differed significantly from spontaneous activity measured in the 50 ms before stimulus presentation (GLM using Poisson distributions and log link function; *p* ≤ 0.05).

Spatially tuned units were then identified using sound-evoked responses collected with the animal at the center of the arena with the head and world coordinate frames in approximate alignment. For a stimulus presentation to be included in this analysis, the animal’s head origin was required to be within 5 cm of a point 2.5 cm behind the arena center (Supplementary Fig. 4). The 2.5 cm offset was applied to provide an approximate account for the distance between the animal’s snout and head center. Head direction was also required to be within ± 15° of the midline of the arena (i.e. the line of symmetry of the arena, so that the animal was facing forward). Sound-evoked responses under these constraints were then fitted with a GLM (Poisson distribution; log link function) with sound source angle relative to the head binned in 30° intervals as predictor. To ensure adequate data for statistical testing, units were only assessed if responses were recorded for ≥ 5 stimulus presentations in each angular bin (186/268 units). Units for which sound source angle significantly reduced model deviance (χ^2^distribution, *p* ≤ 0.05) were classed as spatially tuned (92/186 units). While this approach may not identify all spatially informative neurons (some of which may signal sound location by spike timing rather than rate [18, 54] or that may be tuned only to sounds behind the head that were not sampled by speakers in the aligned condition) it identified a sub population of spatially sensitive units on which further analysis could be performed.

To calculate spatial tuning curves, analysis was expanded to include all head positions and directions recorded. To calculate world-based tuning curves, mean firing rate across trials (0 - 50 ms) was plotted for each sound source angle relative to the arena. For head-based tuning curves, sound source angle relative to the head was binned at 30° intervals and mean spike rate plotted as a function of the bin center angle. To study super-resolution tuning of egocentric units (Fig 5a-b), the bin width used to calculate curves was reduced to 20°, 10°, 5°, 2°, or 1°. To compare spatial tuning of egocentric units with other studies, we also calculated preferred location, modulation depth and tuning width and equivalent rectangular receptive field width for spatial tuning curves calculated relative to the head across 360° according to the methods used for awake cats [15, 18]. For allocentric units, we calculated preferred location and modulation depth for across sound location in the world.

### Modulation depth analysis

For each unit we calculated the depth of spatial modulation for tuning curves in each coordinate frame. Unless otherwise stated, modulation depth (MD) was calculated as:

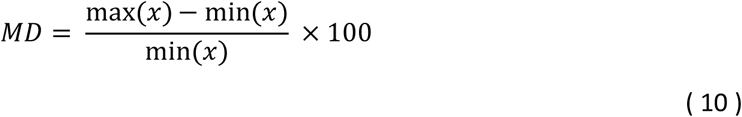

Where *x* is the vector of firing rates in response to sounds located in each 30° bin between ±90° either of the world or head coordinate frame.

Modulation depth could also be calculated for simulated neurons using the same equation but with *x* being a vector of spike probabilities. This approach allowed us to calculate modulation depth for simulated allocentric and egocentric units when presented with sounds during observed animal movement (Fig 3). In simulations, modulation depth could be calculated in head and world coordinate frames that were either represented or irrelevant for neural activity depending on whether the simulation was allocentric or egocentric. We termed modulation depth in the irrelevant coordinate frame *residual modulation* when expressed as a percentage of modulation in the represented coordinate frame:

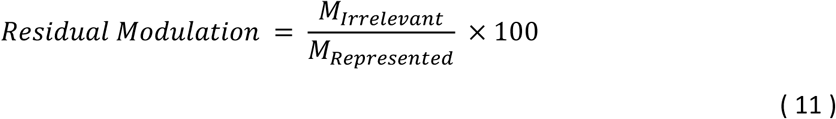

For allocentric simulations, the world was represented and the head was irrelevant; whereas for egocentric simulations, the head was represented and the world was irrelevant.

For each test session in which we observed animal behavior, we compared the relationship between the residual modulation calculated during simulations of both allocentric and egocentric units, with the standard deviation of the head directions (Fig 3). We fitted a linear regression model to this relationship that was subsequently used to test if observed modulation depth values of real units were significantly greater than the residual modulation expected from the animal’s behavior. The linear regression model was fitted using the *fitlm* function in Matlab (R2015a). For each observed unit, we measured the standard deviation of head angles during neural testing (σ) that was used together with the regression models to predict the 95% confidence interval of residual modulation values in the head and in the world coordinate frame. Prediction was performed in Matlab using the *predict* function with the most conservative options selected (simultaneous confidence bounds and prediction for new observations rather than fitted mean values) to give the widest confidence intervals and thus minimize the probability of false positives. If the observed modulation depth of a unit in a particular coordinate frame exceeded the upper bound of the confidence interval for that frame, we identified it as significantly modulated.

### General Linear Models (GLMs)

To compare the relationship between single trial firing rates and sound source angles in the head and world coordinate frames, we fitted the average firing rate on each trial (0-50 ms) with a generalized linear regression model (Matlab, *fitglm* function: Poisson distribution, log link function). For both sound source angles relative to the head and relative to the world, we measured the deviance of models fitted separately with each parameter (D_Test_). The Akaike information criterion was used to compare test models and distinguish allocentric and egocentric units as those for which sound source angle relative to the world or head respectively provided the best model. For all but one unit that was excluded from further analysis, either sound source angle relative to the world or head improved model fit compared to a constant model (analysis of deviance; Bonferroni correction for multiple comparisons; *p* < 0.05).

To visualize GLM performance (Fig 6), we calculated *model fit* for each unit and coordinate frame as:

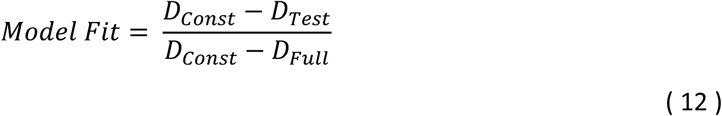

Where D_const_ was the deviance resulting from a constant model, and D_FuM_ was the deviance resulting from a full linear model that included both sound source angle relative to the head and relative to the world. We compared the model fit for data obtained when the head and world coordinate frames were free to vary, and when we restricted data to cases when the head and world coordinate frames were aligned (see above). We also repeated our analysis but with speaker identity or head direction information randomly shuffled between stimulus presentations prior to calculation of spatial tuning curves.

For each analysis in which we calculated model fit, we also calculated *model preference* as:

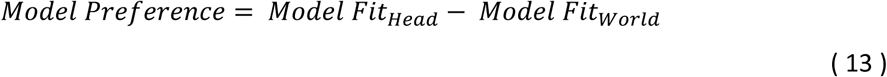

Model preference could thus vary between −100% (better fit for neural data based on sound angle in the world) and +100% (better fit for neural data based on sound angle relative to the head).

For time-based comparison of model performance, we reduced the time over which firing rates were considered (from 50 to 20 ms) and repeated the analysis with a window offset by −60 ms to 90 ms after stimulus presentation that moved with a 2 ms interval. For each unit, we compared the difference between normalized deviance of test models using sound angle relative to the head and relative to the world. Model preference for allocentric and egocentric units were compared using a non-parametric cluster-based statistical test [24] implemented in Matlab through the FieldTrip toolbox [55].

## Acknowledgements

This work was funded by grants to JKB from the BBSRC (BB/H016813/1), the Wellcome Trust / Royal Society (WT098418MA), and Human Frontiers Science Foundation award (RGY0068). OB is funded by the MRC. We would like to thank Prof Jan Schnupp, Prof David McAlpine, Dr Daniel Bendor and Dr Peter Keating for discussions of this work.

